# The making of an olfactory specialist

**DOI:** 10.1101/546507

**Authors:** Thomas O. Auer, Mohammed A. Khallaf, Ana F. Silbering, Giovanna Zappia, Kaitlyn Ellis, Bill S. Hansson, Gregory S.X.E. Jefferis, Sophie Caron, Markus Knaden, Richard Benton

**Author notes:** Corresponding authors, E.

## Abstract

The evolution of animal behaviour is poorly understood. Despite numerous correlations of behavioural and nervous system divergence, demonstration of the genetic basis of interspecific behavioural differences remains rare. Here we develop a novel neurogenetic model, *Drosophila sechellia*, a close cousin of *D. melanogaster* that displays profound behavioural changes linked to its extreme host fruit specialisation. Through calcium imaging, we identify olfactory pathways detecting host volatiles. Mutational analysis indicates roles for individual receptors in long- and short-range attraction. Cross-species allele transfer demonstrates that differential tuning of one receptor is important for species-specific behaviour. We identify the molecular determinants of this functional change, and characterise their behavioural significance and evolutionary origin. Circuit tracing reveals that receptor adaptations are accompanied by increased sensory pooling onto interneurons and novel central projection patterns. This work links molecular and neuronal changes to behavioural divergence and defines a powerful model for investigating nervous system evolution and speciation.

## Introduction

Animals’ perception and responses to the external world adapt, through evolution, to their ecological niche. Despite the incredible phenotypic diversity observed in nature, the genetic and neural basis of behavioural evolution is largely unknown^1,2^. Are there “nodes” in the nervous system that are more susceptible to changes over evolutionary timescales and do they change structurally and/or functionally? What are the underlying molecular mechanisms of evolutionary changes, and do these reflect selection on standing genetic variants or the occurrence of new mutations? Do similar selection pressures on different species lead to the same genetic adaptations?

Some insights have been gained from studies of intraspecific differences in traditional genetic model organisms, such as anxiety behaviours in *Mus musculus*^3^, foraging in *Drosophila melanogaster*^4^, and exploration/exploitation decisions in *Caenorhabditis elegans*^5^. Interspecific behavioural differences are often more dramatic: species within the *Peromyscus* genus of deer mice display variations in many natural behaviours, such as burrowing and parental care^6,7^, while in Nematoda, the predatory *Pristionchus pacificus* exhibits distinct feeding behaviours to *C. elegans*^8^. However, pinpointing the underlying molecular basis of interspecific differences is challenging as it requires that the species are sufficiently comparable molecularly and anatomically, and accessible to genetic manipulations.

Drosophilid flies have emerged as attractive models to define the genetic basis of behavioural evolution: *D. melanogaster* offers a deep foundation of neurobiological knowledge in a numerically-simple brain, and drosophilid species show distinct behavioural traits linked to their diverse ecologies^9^. For example, odour-evoked behaviours of various drosophilids have been correlated with changes in the function of specific sensory pathways^10–15^ and differences in physiological properties of central neurons have been linked to species-specific courtship behaviours^16,17^.

One notable drosophilid is *D. sechellia*, a species endemic to the Seychelles tropical archipelago, which has a common ancestor with the cosmopolitan, ecological generalists *D. melanogaster* and *D. simulans* ∼3 and 0.1-0.24 million years ago, respectively^18–20^ (Fig. 1a). Within this relatively short timespan, *D. sechellia* has evolved extreme specialism for the ripe “noni” fruit of the *Morinda citrifolia* shrub for feeding and oviposition^21,22^. This ecological specialisation is suspected to alleviate interspecific competition, because ripe noni is toxic for other drosophilids^23^; noni may also provide *D. sechellia* with a nutritional source of dopamine precursors to compensate for defects in dopamine metabolism^24^.

**Figure 1.**
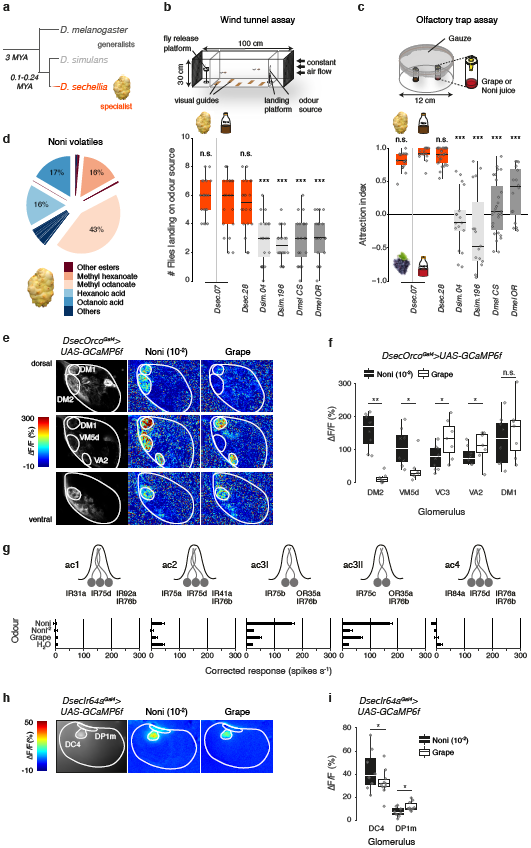
Olfactory behavioural and physiological responses of *D. sechellia* to noni. **a**, *D. sechellia* specialises on noni fruit (*Morinda citrifolia*) while its close cousins *simulans* and *D. melanogaster* are food generalists (MYA = million years ago). **b**, Olfactory responses to noni fruit or noni juice in a wind tunnel assay (schematised at the top) of *D. sechellia* (Drosophila Species Stock Center [DSSC] 14021-0248.07, 14021-0248.28), *D. simulans* (DSSC 14021-0251.004, 14021-0251.196) and *D. melanogaster* (Canton-S and Oregon-R) (n = 20, 10 females/experiment). In this and other panels, boxplots show the median, first and third quartile of the data. Comparisons to *Dsec.07* responses to noni juice are shown in the figure (Kruskal-Wallis test with Dunn’s post-hoc correction): *** *P* < 0.001; n.s. *P* > 0.05. **c**, Olfactory responses in a trap assay (schematised at the top) testing preferences between noni and grape fruit, or between noni and grape juice, for the same drosophilid strains as in **b** (n = 15-27 traps, 22-25 females/trap). Comparisons to *Dsec.07* responses to noni juice are shown in the figure (pairwise Wilcoxon rank-sum test and *P* values adjusted for multiple comparisons using the Benjamini and Hochberg method): *** *P* < 0.001; n.s. *P* > 0.05. **d**, Composition of the odour bouquet of a ripe noni fruit determined by gas chromatography/mass spectrometry (see Methods, Supplementary Fig. 1 and Supplementary Table 1). **e**, Representative odour-evoked calcium responses in the axon termini of Orco OSNs in the *D. sechellia* antennal lobe (genotype: *UAS-GCaMP6f/UAS-GCaMP6f;;DsecOrco*^*Gal*^*4*^^*/+*) acquired by two-photon imaging. Three focal planes are shown, revealing different glomeruli (outlined) along the dorsoventral axis. Left column: raw fluorescence images. Two right columns: relative increase in GCaMP6f fluorescence (ΔF/F%) after stimulation with noni juice (10^−2^) or grape juice. Glomerular identity was defined by position and responses to a panel of diagnostic odours, based upon properties of homologous glomeruli in *D. melanogaster* (Supplementary Fig. 3). **f**, Quantification of odour-evoked calcium responses for the animals represented in **e**. Maximum calcium response amplitudes for each experiment are plotted. Responses to noni (10^−2^) and grape juice were compared using the Wilcoxon signed-rank test: ** *P* < 0.01; * *P* < 0.05; n.s. *P* > 0.05. (n = 7-10, females). **g**, Electrophysiological responses in the antennal coeloconic (ac) sensilla classes to the indicated stimuli (mean ± SEM; n = 6-11, females) in *D. sechellia* (DSSC 14021-0248.07) representing the summed, solvent-corrected activities of the neurons they house, as indicated in the cartoons above. Spike counts for all experiments are provided in Supplementary Table 7. **h**, Representative odour-evoked calcium responses in the axon termini of Ir64a OSNs in the *D. sechellia* antennal lobe (genotype: *UAS-GCaMP6f/UAS-GCaMP6f;;DsecIr64a*^*Gal4*^*/+*) acquired by widefield imaging. Left: raw fluorescence image. Right images: relative increase in GCaMP6f fluorescence (ΔF/F%) after stimulation with noni juice (10^−2^) or grape juice. **i**, Quantification of odour-evoked calcium responses for the animals represented in **h**. Maximum calcium response amplitudes for each experiment are plotted (n = 7-10, females). Responses to noni (10^−2^) and grape juice were compared using the Wilcoxon signed-rank test: ** *P* < 0.01; * *P* < 0.05; n.s. *P* > 0.05.

Concordant with its unique niche, *D. sechellia* exhibits many behavioural differences compared to its close cousins, including olfaction^13–15,23,25^, gustation^26^, courtship and mate choice^27,28^, oviposition^29,30^ and pupariation site selection^31^; several other aspects of its biology have changed, such as increased metabolic tolerance to noni fruit toxins^23,32^. Unbiased mapping approaches have located causal loci for some traits within several – typically large – genomic regions^28,29,31,33^, while candidate approaches have correlated certain phenotypes with changes in the properties of peripheral sensory pathways^13,15,26^.

Despite the potential of *D. sechellia* as a model for comparative neuroscience, investigation of the neural and molecular basis of *D. sechellia*’s behaviours has been hampered by the lack of genetic tools. Here we develop *D. sechellia* into a novel model experimental system, which allows us to move from simple phenotypic correlations to explicitly test the role of specific genetic changes in behavioural evolution.

### Long- and short-range olfactory attraction of *D. sechellia* to noni

In nature, noni-derived volatiles are likely to be the initial cues that guide *D. sechellia* towards its host fruit: field studies demonstrate that these flies can locate noni fruit placed at >50 m^23^. We established two laboratory assays to compare attraction of different wild-type strains of *D. sechellia, D. simulans* and *D. melanogaster* to noni at distinct spatial scales (Fig. 1b-c). To ensure a reproducible odour stimulus, we used commercial noni juice in most experiments. In a long-range, wind tunnel assay^34^, *D. sechellia* displayed significantly higher attraction to the noni source than its sister species (Fig. 1b). Such species-specific attraction was even more pronounced in a short-range olfactory trap assay^15^, where flies were presented with a choice of noni and grape juice (Fig. 1c). In both assays, the level of attraction of *D. sechellia* to noni juice was comparable to that for ripe fruit sources (Fig. 1b-c). This is consistent with the qualitatively similar odour bouquet emitted by juice and ripe fruit, which are dominated by hexanoic and octanoic acids and their methyl ester derivatives (Fig. 1d, Supplementary Fig. 1a-b and Supplementary Table 1).

### Identification of noni-sensing olfactory pathways

Drosophilids detect odours by olfactory sensory neurons (OSNs) housed within morphologically-diverse sensilla on the surface of the antennae and maxillary palps^35^. Most individual OSNs express a single member of the Odorant Receptor (OR) or Ionotropic Receptor (IR) repertoires – which define odour-tuning properties – along with a broadly-expressed co-receptor^36–38^. Neurons expressing the same tuning receptor converge onto a discrete, spatially-stereotyped glomerulus within the primary olfactory centre (antennal lobe) in the brain^35^. Electrophysiological analyses of OSNs in a subset of sensilla in *D. sechellia* have identified several populations that display odour-evoked responses to individual noni volatiles^13–15,39^.

To broadly survey the olfactory representation of the noni bouquet in *D. sechellia*, we generated a transgenic strain in which a *UAS-GCaMP6f* neural activity reporter^40^ was expressed in the majority of OSN populations under the control of Gal4 inserted at the *Or co-receptor* (*Orco*) locus (Supplementary Fig. 2a-d). Two-photon calcium imaging in OSN axon termini in the antennal lobe in *DsecOrco>GCaMP6f* animals revealed a sparse pattern of glomerular activation in response to stimulation of the antenna with noni juice volatiles (Fig. 1e). The global anatomical conservation of the *D. sechellia* antennal lobe to that of *D. melanogaster* and the use of diagnostic odours for specific OSN populations (Supplementary Fig. 3a) allowed us to identify five glomeruli (DM2, VM5d, VC3, VA2, DM1) – out of 35 labelled by *Orco-Gal4* (Supplementary Fig. 2d) – that displayed reliable responses to noni odours (Fig. 1e-f). Of these, only two (DM2 and VM5d) were distinguished by their very high sensitivity to noni juice compared to grape juice (Fig. 1e-f). These glomeruli are innervated by OSNs housed in a common antennal basiconic sensillum class, ab3, which was previously shown to respond to individual noni odours by electrophysiological analysis^13,14^.

To monitor activity in the more limited number of IR-expressing OSNs, we surveyed noni juice sensitivity by electrophysiological recordings in the antennal coeloconic (ac) sensilla classes. Two of these, ac3I and ac3II, responded strongly to noni (Fig. 1g), consistent with analyses using individual noni volatiles (Supplementary Fig. 3b)^15^. One population of Ir OSNs, expressing IR64a, is housed in the sacculus, an internal pocket inaccessible to peripheral electrophysiological analysis^41^. We therefore generated *DsecIr64a>GCaMP6f* animals (Supplementary Fig. 2), which allowed us to measure responses in the two corresponding glomeruli, DP1m and DC4. DC4 neurons were activated slightly more by noni than grape juice, while those innervating DP1m displayed marginally higher responses to grape juice (Fig. 1h-i).

Together these experiments suggest that sensitive detection of noni is mediated by a relatively small set of olfactory pathways in *D. sechellia*.

### Genetic identification of noni-sensing olfactory receptors

To further characterise these noni-sensing olfactory channels, we determined their response profile to a set of prominent individual noni odours (Fig. 1d and Supplementary Fig. 1) using single sensillum recordings, and mutated candidate olfactory receptors by CRISPR/Cas9-mediated genome editing (Supplementary Fig. 4-5). In wild-type ab3 sensilla, the two OSNs can be distinguished by spike amplitude: the larger-spiking ab3A neuron responds most strongly to several methyl esters and only weakly to other odours, while the smaller-spiking ab3B neuron responds strongly to *2*-heptanone and *1*-hexanol (Fig. 2a). In *DsecOrco* mutants, all of these responses are lost (Fig. 2a), consistent with their dependence upon OR signalling.

**Figure 2.**
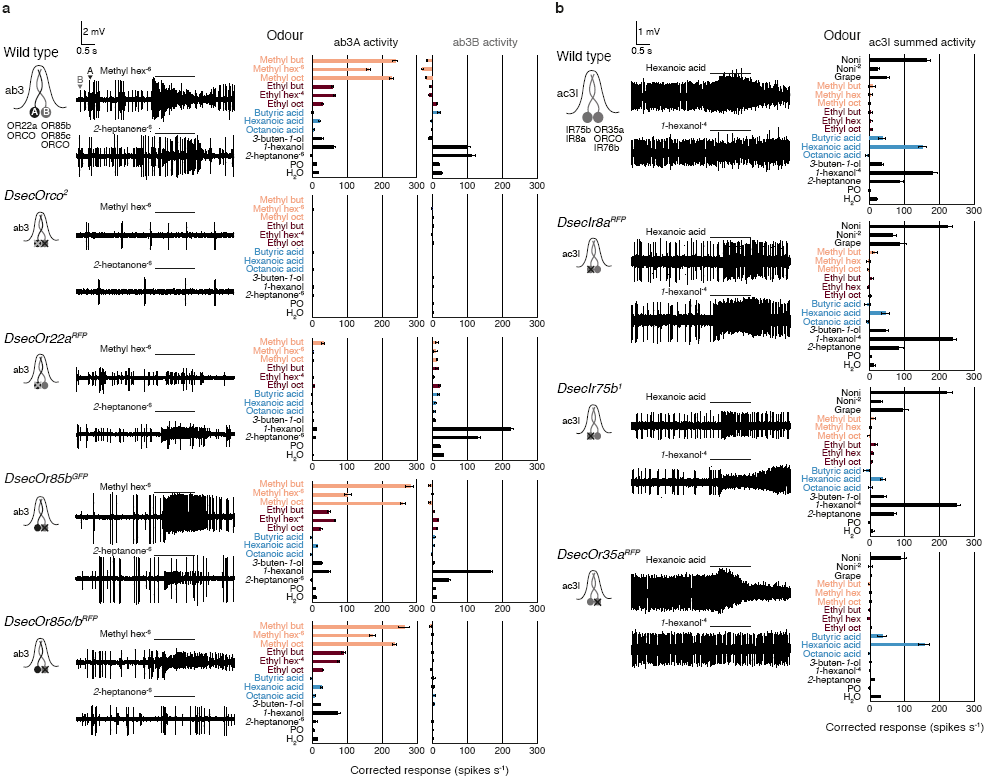
Genetic identification of noni-sensing *D. sechellia* olfactory receptors. **a**, Electrophysiological responses in the two neurons of the ab3 sensillum (indicated in the cartoon on the left) to odours present in noni (mean ± SEM; n = 5-11, females) in wild-type *D. sechellia* (DSSC 14021-0248.07) and olfactory receptor mutants affecting the ab3A neuron (*DsecOr22a*^*RFP*^), ab3B neuron (*DsecOr85b*^*GFP*^, *DsecOr85c/b*^*RFP*^) or both (*DsecOrco*^*^2^*^). Representative response traces to methyl hexanoate (10^−6^) and *2*-heptanone (10^−6^) are shown to the left. Histograms represent the solvent-corrected activities of individual neurons (distinguished by spike amplitude, see arrowheads in wild-type traces). For noni juice, reliable assignment of activity to individual neurons was not possible (data not shown). Odours here and in subsequent figures are coloured according to chemical class: methyl esters (salmon), ethyl esters (dark red), acids (light blue), others (black). Oct = octanoate, hex = hexanoate, but = butanoate. Odours were used at a concentration of 10^−2^ v/v, unless indicated otherwise. We used a lower concentration of some odours as higher doses evoked extremely high firing frequencies accompanied by rapid “pinching” of spike train amplitudes, which precluded accurate quantification. **b**, Electrophysiological responses in the ac3I sensillum (neurons housed are indicated in the cartoon on the left) to noni juice, grape juice and odours present in noni (mean ± SEM; n = 5-10, females only) in *D. sechellia* (DSSC 14021-0248.07) and olfactory receptor mutants affecting either the Ir75b (*DsecIr8a*^*RFP*^, *DsecIr75b*^*^1^*^) or Or35a neuron (*DsecOr35a*^*RFP*^). Representative response traces to hexanoic acid (10^−2^) and *1*-hexanol (10^−4^) are shown to the left. Histograms represent the summed, solvent-corrected activities of both neurons in this sensillum (which cannot be confidently distinguished by spike amplitude). Note that the Or35a expressing neuron shows some residual responses to hexanoic acid in the *DsecIr8a*^*RFP*^ and *DsecIr75b*^*^1^*^ olfactory receptor mutants.

In *D. melanogaster*, ab3A expresses the tandem *Or22a+Or22b* receptor genes^42^. *D. sechellia* possesses only an intact *Or22a* locus^13^, and targeted mutation of this receptor abolishes odour-evoked responses of ab3A, but not ab3B (Fig. 2a). *D. melanogaster* ab3B expresses *Or85b*^35,43^. Unexpectedly, *DsecOr85b* mutants retain some sensitivity to noni odours in these neurons (Fig. 2a), indicating the presence of a redundant receptor. We hypothesised that this is encoded by the neighbouring, closely-related *Or85c* gene: although this locus was reported to be larval-specific in *D. melanogaster*^43,44^, transcripts for this gene are detected in the *D*. *sechellia* adult antennal transcriptome^45^. Consistent with this hypothesis, simultaneous mutation of *DsecOr85b* and *DsecOr85c* led to a complete loss of responses in ab3B, without affecting ab3A (Fig. 2a).

We previously showed that *D. sechellia* ac3I/II sensilla house two OSNs, which express *Or35a* and either *Ir75b* (for ac3I) or *Ir75c* (for ac3II)^15^. We focused on the ac3I class, as this has evolved novel sensitivity to hexanoic acid in *D. sechellia* due to changes in IR75b^15^. Although OSN spike amplitudes could not be reliably distinguished in ac3I (Fig. 2b), we found that mutations in *DsecIr8a* – which encodes the co-receptor for acid-sensing IRs^37^ – or *DsecIr75b* led to selective loss of hexanoic (and butyric) acid responses (Fig. 2b and Supplementary Fig. 5). Conversely, mutation of *DsecOr35a* diminished responses to all other odours except these acids, in-line with the broad tuning of this receptor in *D. melanogaster*^46^.

### Individual ORs are required for long-range attraction

The generation of olfactory receptor mutants affecting the detection of specific noni volatiles allowed us to assess their contribution to the attraction behaviours of *D. sechellia* towards noni. In the long-range olfactory assay, *DsecOrco* mutants – which presumably lack all OR-dependent olfactory pathways – exhibit no attraction to the odour source, with almost all flies failing to reach the landing platform (Fig. 3a). Strikingly, both *DsecOr22a* and *DsecOr85c/b* mutants display similarly strong defects (Fig. 3a). By contrast, *DsecOr35a* mutants were not impaired in this assay (Fig. 3a). Loss of IR8a also led to a significant decrease in long-range attraction in *D. sechellia* (Fig. 3a). This does not appear to be primarily due to defects in the hexanoic acid-sensing pathway, as *DsecIr75b* mutants had either no or milder defects than *DsecIr8a* mutants (Fig. 3a). Mutations in *DsecIr64a*, which encodes (together with IR8a) a broadly-tuned volatile acid sensor in *D. melanogaster*^41,47^ had no effect on this behaviour.

**Figure 3.**
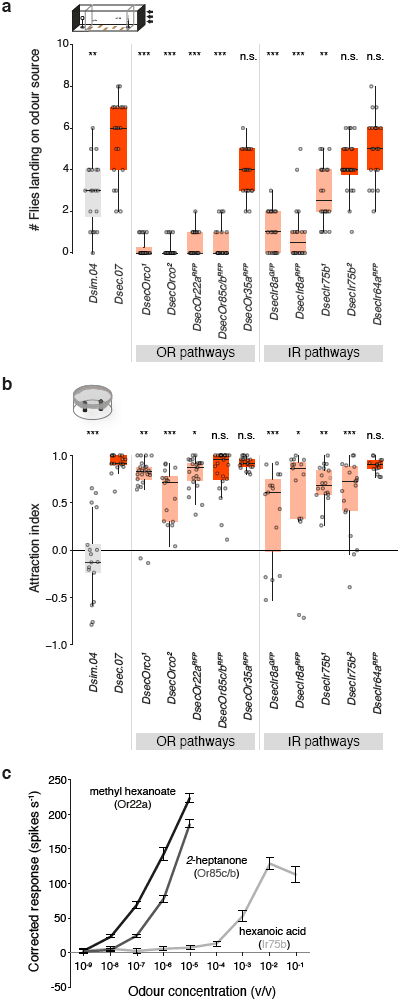
Short- and long-range attractive behaviours of *D. sechellia* require different olfactory pathways. **a**, Olfactory responses to noni juice in the wind tunnel assay in the indicated genotypes (n = 20, 10 females/experiment). The data for *Dsim.04* and *Dsec.07* is repeated from Fig. 1b. Comparisons to *Dsec.07* responses are shown (Kruskal-Wallis test with Dunn’s post-hoc correction): red bars = *D. sechellia* flies with no significant difference to *Dsec.07*; salmon bars = significantly different response; *** *P* < 0.001; ** *P* < 0.01; * *P* < 0.05; n.s. *P* > 0.05. **b**, Olfactory responses in a trap assay testing preference of the indicated genotypes for noni juice or grape juice (n = 15-23 traps, 22-25 females/trap). The data for *Dsim.04* and *Dsec.07* is repeated from Fig. 1c. Comparisons to *Dsec.07* responses are shown (pairwise Wilcoxon rank-sum test and *P* values adjusted for multiple comparisons using the Benjamini and Hochberg method): red bars = *D. sechellia* flies with no significant difference to *Dsec.07*; salmon bars = significantly different response; *** *P* < 0.001; ** *P* < 0.01; n.s. *P* > 0.05. **c**, Electrophysiological responses of wild-type Or22a, Or85c/b and Ir75b neurons in *D. sechellia* upon stimulation with increasing concentrations of their best odour agonists (mean ± SEM, n = 7-15, females). The contribution of Or35a neurons (whose spiking is difficult to separate from Ir75b neurons in ac3I) to hexanoic acid responses is likely to be minimal (Fig. 2b).

In the short-range olfactory assay, *DsecOrco* mutant flies displayed reduced but not abolished, attraction to noni (Fig. 3b). In contrast to the long-range assay, individual Or pathway mutants had very slight (*DsecOr22a*) or no (*DsecOr85c/b, DsecOr35a*) influence on this behaviour (Fig. 3b). *DsecIr8a* mutants displayed reduced attraction, with notable frequent reversals in olfactory preference in several trials (Fig. 3b). A similar phenotype was seen in mutants of *DsecIr75b*, but not *DsecIr64a* (Fig. 3b).

Together these results reveal a critical role for the Or22a and Or85c/b olfactory channels in long-range, but not short-range, attraction of *D. sechellia* to noni. By contrast, short-range attraction evidently depends upon multiple, partially redundant sensory inputs, including the Ir75b pathway.

The different spatial scales at which these olfactory channels contribute to attraction behaviour may be related to their physiological properties: Or22a and Or85c/b neurons have a detection threshold for their best agonists (methyl hexanoate and *2*-heptanone, respectively) that is >1000-fold lower than that of Ir75b neurons for hexanoic acid (Fig. 3c). In addition, the higher volatility of the OR agonists (methyl hexanoate: 3.95 mm/Hg; *2*-heptanone: 3.85 mm/Hg; hexanoic acid: 0.04 mm/Hg; all at 25°C) may result in their more distant diffusion from the odour source in nature.

### Tuning of OR22a is important for species-specific behaviour

Given the crucial role of OR22a and OR85c/b in long-range attraction, we focussed on how these sensory pathways have changed in *D. sechellia* compared to *D. melanogaster* and *D. simulans*. Dose response curves of Or85c/b neurons to their best noni agonist, *2*-heptanone (Supplementary Fig. 6a), revealed an indistinguishable sensitivity across species (Fig. 4a). By contrast, both *D. sechellia* and *D. simulans* Or22a neurons exhibit increased physiological sensitivity to their best agonist, methyl hexanoate, compared to *D. melanogaster* (Fig. 4a and Supplementary Fig. 6a). This result extends a previous comparison of *D. sechellia* and *D. melanogaster*^13,39^, and suggests that increased sensitivity to this odour must already have existed in the last common ancestor of *D. sechellia* and *D. simulans*.

**Figure 4.**
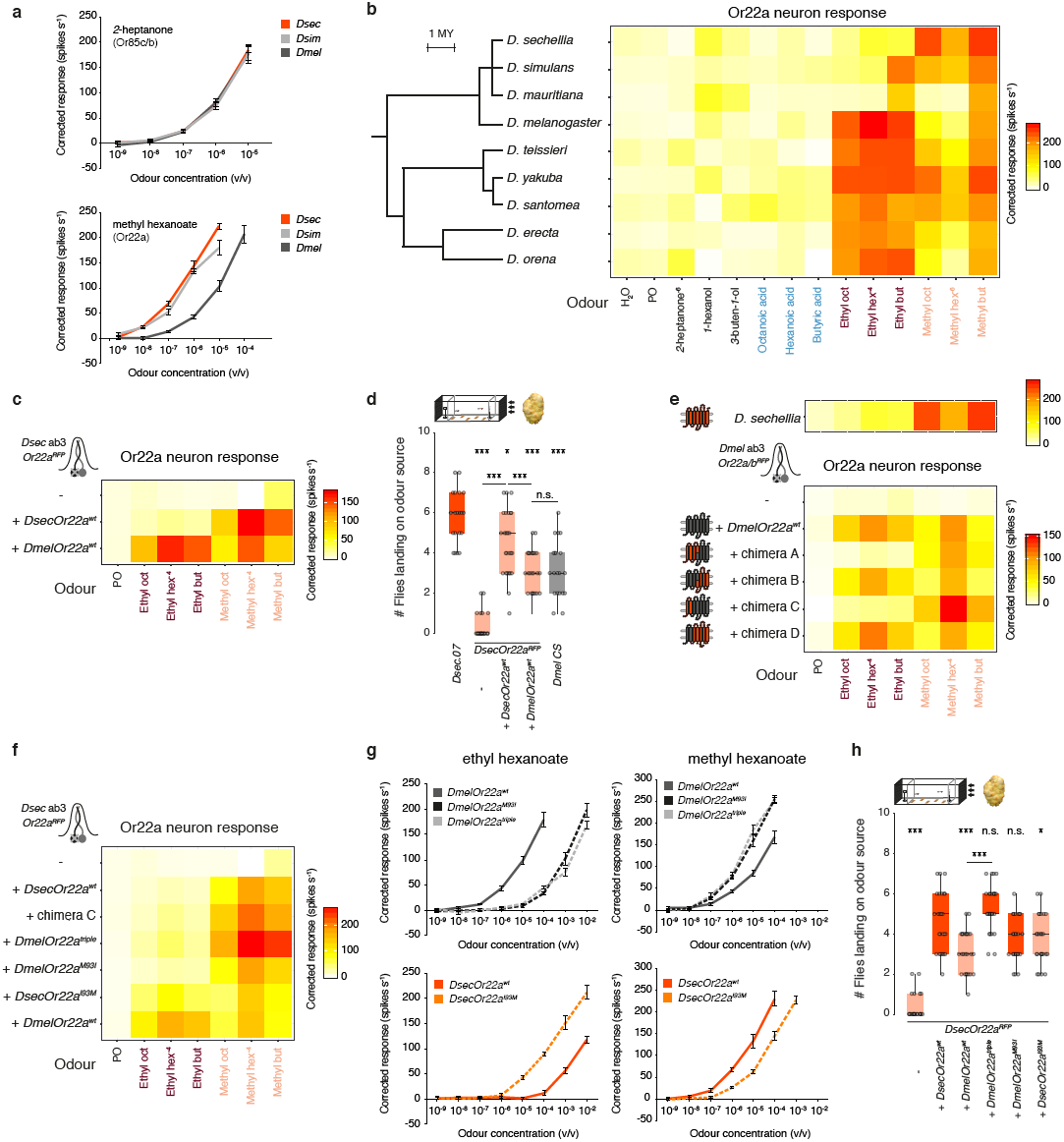
Tuning of OR22a is important for species-specific noni attraction. **a**, Electrophysiological responses of Or85c/b neurons (top) and Or22a neurons (bottom) in the indicated species (*D. sechellia* DSSC 14021-0248.07, *D. simulans* DSSC 14021-0251.004, *D. melanogaster* Canton-S) upon stimulation with increasing concentrations of *2*-heptanone and methyl hexanoate, respectively, (mean ± SEM, n = 9-11, females). **b**, Electrophysiological responses of Or22a neurons to noni odours across the *D. melanogaster* species subgroup: *D. sechellia* (DSSC 14021-0248.07), *D. simulans* (DSSC 14021-0251.004), *D. mauritiana* (DSSC 14021-0241.151), *D.melanogaster* (Canton-S), *D. yakuba* (DSSC 14021-0261.00), *D. santomea* (DSSC 14021-0271.00), *D. teissieri* (DSSC 14021-0257.01), *D. orena* (DSSC 14021-0245.01), and *D. erecta* (DSSC 14021-0224.01); MY = million years. The colour scale bar for averaged, solvent-corrected spike counts s^−1^ is shown on the far-right (n = 5-11, females). **c**, Physiological responses of OR22a protein orthologues of *D. melanogaster* and *D. sechellia* integrated at the *Or22a* locus of *D. sechellia* (n = 6-11, females). **d**, Olfactory responses to noni fruit in the wind tunnel assay of the indicated genotypes (n = 20, 10 females/experiment). Comparisons to *Dsec.07* responses are shown if not indicated otherwise (Kruskal-Wallis test with Dunn’s post-hoc correction): salmon bars indicate *D. sechellia* genotypes with significantly different response to *Dsec.07*; *** *P* < 0.001; ** *P* < 0.01; * *P* < 0.05. **e**, Physiological responses of chimeric OR22a proteins integrated at the *Or22a/b* locus of *D. melanogaster* (n = 5-6, females, see also Supplementary Fig. 6b) towards a panel of noni odours. Upper row: *D. sechellia* wild-type response as shown in **b**. Schematics on the left indicate the relative proportions of *D. sechellia* (red) and *D. melanogaster* (dark grey) sequence to each chimera (compare Supplementary Fig. 6c for details). **f**, Physiological responses of OR22a protein variants integrated at the *Or22a* locus of *D. sechellia* (n = 6-11, females). **g**, Left column: electrophysiological responses of *D. sechellia* Or22a neurons expressing the indicated transgenes upon stimulation with increasing concentrations of ethyl hexanoate. Right column: electrophysiological responses of *D. sechellia* Or22a neurons expressing the indicated transgenes upon stimulation with increasing concentrations of methyl hexanoate (mean ± SEM, n = 10, females). **h**, Olfactory responses to noni fruit in the wind tunnel assay of the indicated genotypes (n = 20, 10 females/experiment). Comparisons to *DsecOr22a* mutants rescued with *DsecOr22a*^*wt*^ cDNA responses are shown if not indicated otherwise (Kruskal-Wallis test with Dunn’s post-hoc correction): red bars = *D. sechellia* flies with no significant difference to *DsecOr22a* mutants rescued with *DsecOr22a*^*wt*^ cDNA; salmon bars = significantly different response; *** *P* < 0.001; ** *P* < 0.01; * *P* < 0.05; n.s. *P* > 0.05.

Broader ligand profiling of Or22a neurons in all members of the *D. melanogaster* species subgroup (Fig. 4b) revealed that *D. sechellia* was the only species in which these neurons display selective high sensitivity to methyl esters; others, including *D. simulans*, respond to at least a subset of ethyl esters (Fig. 4b). The specificity of *Dsec*OR22a for methyl esters may be related to the unusually elevated levels of these volatiles over ethyl esters in noni, when compared to other fruits (Supplementary Fig. 1c). Together, these observations suggest that changes in tuning sensitivity and/or breadth of OR22a – but not OR85c/b – in *D. sechellia* contribute to the differential behaviour of this species, and led us to focus our attention on this olfactory pathway.

To assess the significance of these differences for *D. sechellia*’s attraction to noni, we reintroduced either *DsecOr22a*^*wt*^ or *DmelOr22a*^*wt*^ into the *DsecOr22a* mutant background at the endogenous locus (Supplementary Fig. 6b). We tested both the electrophysiological responses conferred by these different receptor alleles and the long-range attraction of the transgenic flies to noni fruit (Fig. 4c-d). *Dsec*OR22a^wt^ can restore higher sensitivity responses to methyl hexanoate (and other methyl esters) than *Dmel*OR22a^wt^, indicating that the receptor itself is a key determinant of species-specific neuron tuning properties (Fig. 4c). Consistent with these physiological differences, *Dsec*OR22a^wt^, but not *Dmel*OR22a^wt^, can rescue the long-range behavioural responses to almost wild-type levels (Fig. 4d). These results indicate that tuning of OR22a can explain, in part, the species-specific odour-driven attraction to noni fruit.

### Molecular basis of tuning changes in OR22a

We next sought the molecular basis of the tuning changes in *Dsec*OR22a. Transgenic expression of chimeric versions of *Dsec*OR22a^wt^ and *Dmel*OR22a^wt^ in Or22a neurons in *D. melanogaster* (replacing the endogenous *Or22a/Or22b* loci; Supplementary Fig. 6b) indicated that high-sensitivity and selectivity for methyl esters versus ethyl esters were determined by the N-terminal 100 amino acids of *Dsec*OR22a (chimera C) (Fig. 4e). Within this region, we focussed on the three positions in *Dmel*OR22a (I45, I67 and M93) in which this receptor differs from the OR22a orthologues in the species displaying narrowed tuning for methyl esters (i.e., *D. sechellia, D. simulans* and *D. mauritiana*) (Fig. 4b and Supplementary Fig. 6c). We generated a site-directed mutant receptor in which these three residues were mutated to those present in the other species (i.e., *Dmel*OR22a^I45V,I67M,M93I^; hereafter, *Dmel*OR22a^triple^). This variant was only responsive to methyl esters, similar to *Dsec*OR22a^wt^ (Supplementary Fig. 6d). Individual mutation of these residues revealed that all impact odour tuning, but in different ways (Supplementary Fig. 6d), with *Dmel*OR22a^M93I^ most faithfully recapitulating the relatively higher sensitivity of this receptor to methyl esters over ethyl esters.

To assess the significance of these molecular differences for *D. sechellia*’s attraction to noni, we reintroduced different wild-type and mutant alleles of *Or22a* into the mutant *DsecOr22a* locus. As observed in the *D. melanogaster* expression system, OR22a chimera C, *Dmel*OR22a^triple^ and *Dmel*OR22a^M93I^ all display narrowed sensitivity to methyl esters (Fig. 4f). Conversely, *Dsec*OR22a^I93M^ exhibits broadened sensitivity to both ester classes (Fig. 4f). Dose-response analysis revealed a marked decrease in sensitivity of *Dmel*OR22a^triple^ and *Dmel*OR22a^M93I^ to ethyl hexanoate, and increased sensitivity of these receptor variants to methyl hexanoate (Fig. 4g). *Dsec*OR22a^I93M^ displayed the opposite changes in sensitivity compared to *Dsec*OR22a^wt^ (Fig. 4g).

In the long-range olfactory behaviour assay, *Dmel*OR22a^triple^ restored *D. sechellia*-like attraction to noni (Fig. 4h), while both *Dmel*OR22a^M93I^ and *Dsec*OR22a^I93M^ displayed attraction levels that are intermediate between those of the two species wild-type receptor rescue (Fig. 4h). These observations provide evidence that these molecular differences in OR22a orthologues contribute to species-specific olfactory behaviours.

### Conserved wiring but divergent sensory representation scales of Or22a

The *Or22a* allele swap experiments indicate an important role for receptor tuning in defining species-specific olfactory attraction. Nevertheless, the observation that *D. simulans* does not display the same high attraction to noni as *D. sechellia* (Fig. 1b-c), despite similar (though not identical) Or22a neuron response properties (Fig. 4a-b), suggests additional changes in *D. sechellia* are important. Previous comparison of *D. melanogaster* and *D. sechellia* revealed an expansion of the ab3 sensillum population^13,39^, which we found is reflected in a nearly three-fold increase in the number of Or22a-expressing neurons (Fig. 5a-b). Importantly, within this species trio, the expansion is restricted to *D. sechellia*, and conserved across different wild-type strains (Fig. 5a-b). The Or85c/b neuron population displays a similar expansion (data not shown), concordant with these neurons being paired with Or22a neurons in ab3 sensilla (Supplementary Fig. 4d). This expansion does not reflect a global increase in antennal OSN numbers (Supplementary Fig. 7a and ^15^); some populations, such as ab1 Or42b neurons, are reduced in size in *D. sechellia* (Fig. 5b).

**Figure 5.**
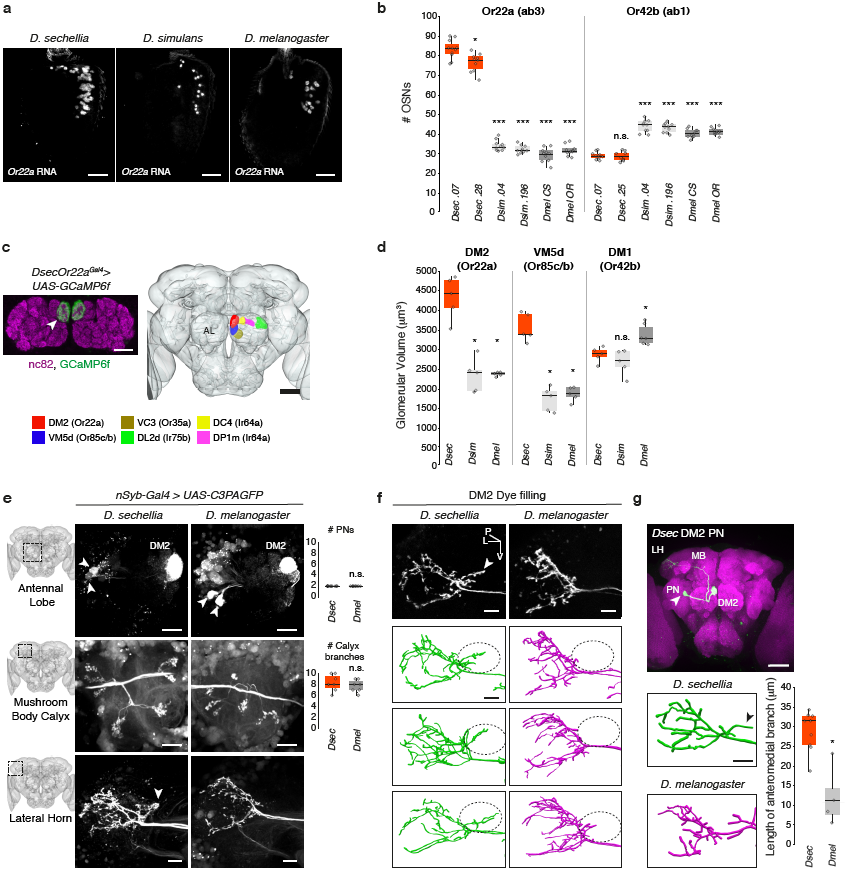
Neuroanatomical analysis of noni-sensing olfactory pathways. **a**, *Or22a* RNA expression in whole-mount antennae of *D. sechellia* (DSSC 14021-0248.07), *D. simulans* (DSSC 14021-0251.004) and *D. melanogaster* (Canton-S). Scale bar = 25 µm. **b**, Quantification of the number of OSNs expressing Or22a (ab3; left) or Or42b (ab1; right) in *D. sechellia* (DSSC 14021-0248.07, DSSC 14021-0248.28, DSSC 14021-0248.25), *D. simulans* (DSSC 14021-0251.004, DSSC 14021-0251.196), and *D. melanogaster* (Canton-S, Oregon-R) (n = 8-11, females). Comparisons to *Dsec.07* cell number counts are shown (pairwise Wilcoxon rank-sum test and *P* values adjusted for multiple comparisons using the Benjamini and Hochberg method): *** *P* < 0.001; * *P* < 0.05; n.s. *P* > 0.05. **c**, Left: expression of *Or22a*^*Gal4*^-driven GCaMP6f signal (detected by α-GFP) in the DM2 glomerulus (arrowhead) of the antennal lobe (the neuropil is visualised with nc82 (magenta); genotype: *UAS-GCaMP6f/+;DsecOr22a*^*Gal4*^*/+;*). Scale bar = 25 µm. Right: Segmentation of antennal lobe glomeruli in the *D. sechellia* brain. Scale bar = 50 µm. **d**, Quantification of glomerular volumes of DM2 (innervated by *Or22a*-expressing neurons), VM5d (*Or85c/b*) and DM1 (*Or42b*) between species (n = 5, females). Comparisons to *D. sechellia* glomerular volumes are shown (pairwise Wilcoxon rank-sum test and *P* values adjusted for multiple comparisons using the Benjamini and Hochberg method): * *P* < 0.05; n.s. *P* > 0.05. **e**, Labelling of DM2 innervating projection neurons (PNs) via photoactivation in *D. sechellia* and *D. melanogaster* (see Methods for details; genotypes *D. sechellia DsecnSypGal4/UAS-C3PAGFP;; D. melanogaster; UAS-SPAGFP/UAS-C3PAGFP;nSybGal4/UAS-C3PAGFP*). Schematics on the left depict the site of image acquisition. Upper panel: antennal lobe with labelled PNs (arrowheads) and DM2 glomerulus. Note that there is some background expression from the GFP transgenes in PNs even without photoactivation, Scale bar = 20 µm; right: quantification of the number of DM2-innervating PNs in *D. sechellia* and *D. melanogaster* (n = 7, females, grey circles) (pairwise Wilcoxon rank-sum test): n.s. *P* > 0.05. Middle panel: PN innervation of the mushroom body calyx, Scale bar = 10 µm; right: quantification of the number of branches innervating the mushroom body calyx in *D. sechellia* and *D. melanogaster* (n = 7, females) (pairwise Wilcoxon rank-sum test): n.s. *P* > 0.05. Lower panel: axonal branches of PNs in the lateral horn. Arrowhead: extra branch in the lateral horn of *D. sechellia*. Scale bar = 10 µm. **f**, Axonal arbours in the lateral horn of dye-filled DM2 PNs in *D. sechellia* and *D. melanogaster*. The arrowhead indicates the extra branch in *D. sechellia*. Below: Tracing of axonal branches in both species for three representative samples. The circle depicts the position of the *D. sechellia*-specific axonal branch. The schematic on the left shows the site of image acquisition in the brain. P = posterior, L = lateral, V = ventral. Scale bar = 10 µm. **g**, A single dye-filled DM2 PN in *D. sechellia*. Below: traced, representative example of a single DM2 innervating PN in *D. sechellia* and *D. melanogaster*. Scale bar = 10 µm. Left: Quantification of PN branch length between species (n = 4-9, females) (pairwise Wilcoxon rank-sum test): * *P* < 0.05.

To analyse the central projections of OSN populations in *D. sechellia*, we inserted Gal4 at the corresponding receptor loci, and combined these with *UAS-GCaMP6f* as a fluorescent (GFP-based) anatomical marker (Supplementary Fig. 2a-b). These tools allowed us to extend observations from single-neuron dye-filling analyses^13–15^, demonstrating that OSN glomerular innervation patterns are indistinguishable between *D. sechellia* and *D. melanogaster* (Fig. 5c and Supplementary Fig. 2c-d). However, concordant with the expansion of the sensory populations, the glomerular targets of Or22a and Or85c/b neurons (DM2 and VM5d, respectively) are nearly doubled in volume in *D. sechellia* compared to *D. melanogaster* and *D. simulans* (Fig. 5d).

### Neuroanatomical differences in a brain centre for innate behaviour

To visualise higher-order elements of the Or22a pathway in *D. sechellia*, we generated a pan-neuronal driver (*nSyb-Gal4*) (Supplementary Fig. 7b) and combined this with a photoactivatable GFP transgene (*UAS-C3PA*)^48^ for targeted photolabelling of projection neurons (PNs) that synapse with Or22a OSNs in DM2 (see Methods). Analysis with analogous reagents in *D. melanogaster* permitted comparison of these species. Two DM2 projection neurons (PNs) were consistently labelled in both *D. sechellia* and *D. melanogaster* (Fig. 5e) indicating that the OSN population expansion is not accompanied by an increase in the number of their postsynaptic partners.

PNs innervate two higher olfactory centres, the mushroom body (required for olfactory memory storage and retrieval) and the lateral horn (a region implicated in innate olfactory responses)^49^. Within the mushroom body, the number and arrangement of PN axonal branches was similar between *D. sechellia* and *D. melanogaster* (Fig. 5e). In the lateral horn, the global anatomy was conserved, with the main tract bifurcating into dorsal and ventral branches. However, we observed a more prominent axonal branch in *D. sechellia* DM2 PNs dorsally of this bifurcation point that innervates an area of the lateral horn not targeted by the labelled homologous *D. melanogaster* neurons (Fig. 5e). Similar observations were made by visualising PNs through targeted electroporation of a lipophilic dye^48^ into the DM2 glomerulus (Fig. 5f). To quantify this difference, we combined photo- and dye-labelling to visualise single DM2 PNs (see Methods) and traced the resultant axonal arbours in the lateral horn. Measurements of the respective axonal arbour after 3D reconstruction confirmed the presence of a longer *D. sechellia*-specific branch extending towards the anterior-medial part of the lateral horn (Fig. 5g).

## Discussion

We have developed *D. sechellia* as a neurobiological model to link genetic and neural circuit changes to behaviours relevant for its striking ecology. Through characterisation of the role of the Or22a sensory pathway in this species and comparison of this circuit’s functional and structural properties across closely-related drosophilids, our use of this model already provides several insights into behavioural evolution.

First, while changes in peripheral sensory abilities are often correlated with species-specific behaviours^2^, the *Or22a* allele transfer experiments in *D. sechellia* provide, to our knowledge, the first direct evidence that peripheral olfactory receptor tuning properties contribute to species-specific odour-evoked behaviour.

Second, our mapping of molecular determinants underlying the re-tuning of OR22a is informative for our understanding of the molecular basis of odour/receptor interactions. When mapped onto a presumably homologous ORCO structure^50^, the key molecular change (OR22a^M93I^) is located within a putative ligand-binding pocket (Supplementary Fig. 8a-b). Furthermore, this residue corresponds to the same site that is linked with ligand-sensitivity differences in a highly divergent receptor, *Dmel*OR59b^51^ (Supplementary Fig. 8c), suggesting that this position is a “hot-spot” for functional evolution. Importantly, the relevant mutations in *Or22a* did not emerge in the *D. sechellia* lineage, but rather were present in its last common ancestor with *D. simulans* and *D. mauritiana*. Additional mutations in the latter two species must have occurred to account for the differences in ethyl ester sensitivity.

Third, although functional differences in this receptor are important, they do not entirely explain *D. sechellia*’s olfactory adaptations, given the similarity in Or22a neuron response profiles of *D. simulans* and *D. sechellia*. The *D. sechellia*- specific expansion of the Or22a neuron population is likely to be a key additional evolutionary innovation. We note that the population expansion alone is insufficient, as shown by the inability of *Dmel*OR22a to restore *D. sechellia*-like host attraction when expressed in this larger population of OSNs. Or22a postsynaptic partner PNs have not changed in number; the consequent increased pooling of sensory signals onto these interneurons could, for example, enhance the signal-to-noise ratio of this sensory transformation. Moreover, the difference in PN projections raises the possibility that central circuit connectivity changes form part of *D. sechellia*’s adaptation to noni fruit. Development of tools in additional strains and species will be necessary to understand the evolutionary significance of this intriguing wiring variation.

Fourth, the profound role we describe for *D. sechellia* OR22a in host attraction may explain the rapid molecular evolution of this locus across species: *Or22a* displays substantial intra- and interspecific nucleotide and copy number variation^8,52–54^, concordant with diversification in physiological responses of presumed *Or22a*-paralog expressing neurons in a variety of drosophilids^11,12,55^. In addition, *D. erecta*, a specialist on *Pandanus* fruit, may also exhibit expansion of this sensory population^11^. Interestingly, a second noni-adapted drosophilid, *D. yakuba mayottensis* was recently identified^56^. However, analysis of this species’ *Or22a* sequence does not reveal the same genetic changes we identified within *DsecOr22a* (Supplementary Fig. 9a). Furthermore, neither the OR22a response profile nor OSN numbers deviate from other *D. yakuba* strains (Supplementary Fig. 9b-c). These observations imply the existence of an independent evolutionary solution to locate a common host fruit.

Finally, although we have focussed on the Or22a pathway, we have shown that several other olfactory channels are important for noni attraction. These include Or85c/b neurons, which have conserved physiological properties while increasing in number in *D. sechellia*, and Ir75b neurons, which have both changed in function and number, while preserving central projections^15^. These observations indicate that different neural pathways may adapt in distinct ways, possibly reflecting different selection pressures and roles in controlling odour-evoked behaviours. Indeed, our genetic uncoupling of long- and short-range olfactory attraction in *D. sechellia* reveals a previously unappreciated facet of sensory coding in the olfactory system. Future development and application of the genetic toolkit in *D. sechellia* promises to offer both fundamental insights into how genes and neurons control behaviour, and how they change to permit evolution of novel species-specific traits.

## Acknowledgements

We thank Y. Bellaïche, B. Deplancke, K. S. Douglas, S. Lavista Llanos, K. O’Connor-Giles, D. Stern, G. Suh, A. Yassin, the Bloomington *Drosophila* Stock Center (NIH P40OD018537) and the Developmental Studies Hybridoma Bank (NICHD of the NIH, University of Iowa) for reagents, B. Prud’homme for instruction on *Drosophila* microinjections; B. Sutcliffe and S. Cachero for advice on the generation of reference brains, I. Alali for technical assistance, J. Simpson for sharing details about the nSyb promoter construct, P. C. Chai for assistance with glomerular identification and R. Alvarez Ocana for sharing *D. simulans* brain samples. We thank I. Rentero Rebollo, J. R. Arguello, J. Sánchez-Alcañiz, L. Prieto-Godino and members of the Benton laboratory for discussions and comments on the manuscript. T.O.A. is supported by a Human Frontier Science Program Long-Term Fellowship (LT000461/2015-L). M.A.K., B.S.H., and M.K. are supported by the Max Planck Society. K.E. and S.C. are supported by a National Institute of Health Award (1 R01 NS 167970) and a Eunice Kennedy Shriver National Institute of Child Health & Human Development Award of the National Institutes of Health (5 T32 HD 007491). Research in R.B.’s laboratory is supported by an ERC Consolidator Grant (615094), the Swiss National Science Foundation and the Fondation Herbette.

## Author Contributions

T.O.A. and R.B. conceived of the project. All authors contributed to experimental design, analysis and interpretation of results. T.O.A. generated all molecular reagents and new drosophilid mutants and transgenic lines; other experimental contributions were as follows: T.O.A. (Fig. 1c,g; Fig. 2a,b; Fig. 3b,c; Fig. 4a-c,e-g; Fig. 5a,b,d,f; Fig. S2; Fig. S3b; Fig. S4; Fig. S5; Fig. S6; Fig. S7a,b; Fig. S9), M.A.K. (Fig. 1b,d; Fig. 3a; Fig. 4d,h; Fig. S1; Table S1), G.Z. (Fig. 1c; Fig. 3b; Fig. 5a; Fig. S2b; Fig. S4c,e,g; Fig. S5b-e; Fig. S7a; Fig. S9c), A.F.S. (Fig. 1e,f,h,i; Fig. S3a), K.E. (Fig. 5e-g), G.S.X.E.J. (Fig. 5c), R.B. (Fig. 5c). T.O.A. and R.B. wrote the paper with input from all other authors.

## Methods

### Volatile collection, gas chromatography and mass spectrometry

Volatiles were collected from 1 ml of fruit juice or 13 g of noni fruit at different ripening stages in capped 15 ml glass vials with poly-tetra-fluoroethylene-lined silicone septa (Sigma, 23242-U). After penetrating the septum of the cap with a Solid Phase Microextraction (SPME) fibre holder, the SPME fibre (grey hub plain; coated with 50/30 µm divinylbenzene/carboxen on polydimethylsiloxane on a StableFlex fibre (Sigma, 57328-U)) was exposed to the headspace of each vial for 30 min at room temperature. The SPME fibre was retracted and immediately inserted into the inset of a Gas Chromatography-mass spectrometry (GC-MS) system (Agilent 7890B fitted with MS 5977A unit) for desorption at 260°C in split mode (split ratio 100:1). The GC was operated with a HP-INNOWax column (Agilent 19091N-133UI). The sample (SPME) was injected at an initial oven temperature of 50°C; this temperature was held for 1 min and gradually increased (3°C min^−1^) to 150°C before holding for 1 min. Subsequently, the temperature was increased (20°C min^−1^) to 260°C and held for 5 min. The MS-transfer-line was held at 260°C, the MS source at 230°C, and the MS quad at 150°C. MS spectra were taken in EI-mode (70 eV) in a 29-350 m/z range. Between different collections, the SPME fibre was conditioned at 270°C for 15 min. All chromatograms were processed using MSD ChemStation F.01.03.2357 software. Volatile compounds were identified using the NIST library and matched to standards of the Max-Planck-Institute for Chemical Ecology library. For quantification, peak areas were measured for 3 replicates for each sample. Vapour pressure values for hexanoic acid, methyl hexanoate and *2*-heptanone were described previously^57,58^ (www.thegoodscentscompany.com/data/rw1008741.html).

### *Drosophila* strains

*Drosophila* stocks were maintained on standard corn flour, yeast and agar medium under a 12?h light:12?h dark cycle at 25°C. For all *D. sechellia* strains, a few g of Formula 4-24® Instant *Drosophila* Medium, Blue (Carolina Biological Supply Company) soaked in noni juice (nu3 GmbH) were added on top of the standard food. Wild-type *Drosophila* strains are described in the corresponding figure legends. The mutant and transgenic lines generated in this study are listed in Supplementary Table 2.

### CRISPR/Cas9-mediated genome engineering

*sgRNA expression vectors:* for expression of single sgRNAs, oligonucleotide pairs (Supplementary Table 3) were annealed and cloned into *BbsI*-digested *pCFD3-dU6-3gRNA* (Addgene #49410), as described^59^. To express multiple sgRNAs from the same vector backbone, oligonucleotide pairs (Supplementary Table 4) were used for PCR and inserted into *pCFD5* (Addgene #73914) via Gibson Assembly, as described^60^.

*Donor vectors for homologous recombination*: to generate an eGFP-expressing donor vector (*pHD-Stinger-attP*), the fluorophore was excised from *pStinger*^61^ with *NcoI*/*HpaI* and used to replace the *DsRed* sequence in *NcoI*/*HpaI*-digested *pHD-DsRed-attP* (Addgene plasmid #51019)^62^. Homology arms (1-1.6 kb) for individual target genes were amplified from *D. sechellia* (DSSC 14021-0248.07) or *D. melanogaster* (Research Resource Identifier Database:Bloomington *Drosophila* Stock Center [RRID:BDSC]_58492) genomic DNA and inserted either into *pHD-DsRed-attP* or *pHD-Stinger-attP* via restriction cloning. Details and oligonucleotide sequences are available from the authors upon request.

*Transgenic source of Cas9*: *pBac(nos-Cas9,3XP3-YFP)* (gift of D. Stern) was integrated into *D. sechellia* (DSSC 14021-0248.07) via *piggyBac* transgenesis. The insertion was mapped to the fourth chromosome using TagMap^63^.

### Transgene construction

Oligonucleotides for each cloning step are listed in Supplementary Table 5.

*attB-nSyb-Gal4,miniW*: 1.9 kb upstream sequence of the neuronal Synaptobrevin (*nSyb*) gene were amplified from *D. sechellia* genomic DNA (DSSC 14021-0248.07) and inserted into *pGal4attB*^64^ via restriction cloning using *NotI* and *KpnI*. *attB-Gal4,3XP3-Stinger*: we first generated an *eGFPnls-SV40* fragment via PCR (using *pHD-Stinger-attP* as template) and fused it to a minimal *attB40* site^65,66^ before insertion into *pCR-Blunt II-TOPO* (Thermo Fisher). We added a *3XP3-Stinger* fragment amplified from *pHD-Stinger* via restriction cloning using *EcoRV* and *SalI*. Subsequently, we placed a *loxP* site downstream of the initial *SV40* sequence via oligonucleotide annealing and *SpeI* and *KpnI* restriction cloning, to produce *pCR-TOPO-loxP-attB40-eGFPnlsSV40 rev-3XP3:Stinger*. We replaced the *eGFPnls-SV40* sequence with an *hsp70-Gal4-SV40* fragment via PCR amplification of the vector backbone and *Gal4* from *pGal4attB*^64^ and subsequent Gibson Assembly resulting in *attB-Gal4,3XP3-Stinger*.

*attB-Or22a*^*wt*^,*3XP3-Stinger*: in the *pCR-TOPO-loxP-attB40eGFPnlsSV40 rev-3XP3-Stinger* plasmid described above, the *eGFPnlsSV40* fragment was flanked by *EcoRV* and *SalI* sites, which were used to integrate either the *D. sechellia* or *D. melanogaster Or22a* ORF+3’UTR after PCR amplification from cDNA. This resulted in *attB-DsecOr22a*^*wt*^,*3XP3-Stinger* or *attB-DmelOr22a*^*wt*^,*3XP3-Stinger*, respectively.

*Or22a chimeras*: chimeric constructs between *D. sechellia* and *D. melanogaster Or22a* were generated by PCR amplification and fusion using the respective species *Or22a* gene templates. After subcloning into *pCR-Blunt II-TOPO* and sequence confirmation, they were integrated into *pCR-TOPO loxP attB40eGFPnlsSV40rev-3XP3-Stinger* via restriction cloning.

*Or22a site-directed mutant constructs:* point mutations were introduced via site directed mutagenesis following standard procedures.

### *Drosophila* microinjections

Transgenesis of *D. sechellia* and *D. melanogaster* was performed in-house following standard protocols (http://gompel.org/methods). For the *D. sechellia* egg-laying agar plates, we replaced grape juice with noni juice and added a few g of Formula 4-24® Instant *Drosophila* Medium, Blue (Carolina Biological Supply Company) soaked in noni juice (nu3 GmbH) on the surface. Embryos were manually selected for the right developmental stage prior to alignment and injection. For *piggyBac* transgenesis, we co-injected *piggyBac* vector (300 ng µl^−1^) and *piggyBac* helper plasmid^67^ (300 ng µl^−1^). For CRISPR/Cas9-mediated homologous recombination, we injected a mix of a sgRNA-encoding construct (150 ng µl^−1^), donor vector (400 ng µl^−1^) and *pHsp70-Cas9* (400 ng µl^−1^) (Addgene #45945)^68^. The DsRed fluorescent marker was destroyed in *DsecnSyb-Gal4* and *DsecUAS-C3PAGFP* via injection of a sgRNA construct targeting *DsRed* (150 ng µl^−1^) and *pHsp70-Cas9* (400 ng µl^−1^). Injections into *Dsecnos-Cas9* were performed mixing the sgRNA construct (150 ng µl^−1^) and donor vector (500 ng µl^−^ 1). Site-directed integration into *attP* sites was achieved by co-injection of an *attB*-containing vector (400 ng µl^−1^) and either *p3xP3-EGFP.vas-int.NLS* (400 ng µl^−1^) (Addgene #60948)^69^ or *pBS130* (encoding phiC31 integrase under control of a heat shock promoter, Addgene #26290)^70^. All concentrations are the final values in the injection mix.

### Wind tunnel assay

Long-range attraction experiments were performed in a wind tunnel as described previously^34^ with a flight arena of 30 cm width, 30 cm height and 90 cm length. The airstream in the tunnel (0.3 m s^−1^) was produced by a fan (Fischbach GmbH, Neunkirchen, Germany), and filtered through an array of four activated charcoal cylinders (14.5 cm diameter x 32.5 cm length; Camfil, Trosa, Sweden). The wind tunnel was maintained within a climate chamber at 25°C and 50–55% relative humidity under white light. Flies were starved for approximately 20 h; to ensure the flight ability of assayed animals, flies were first released into a mesh cage (50 x 50 x 50 cm, maintained at the same conditions as the wind tunnel) and females escaping from the food vial were collected with an aspirator. For each assay, ten 4-6 days old females were released from a plastic tube (with a mesh covering one end; the open end facing the landing platform) fixed horizontally in the centre of the first 5-10 cm of the downwind end of the tunnel. The landing platform was built by a filter paper (3 x 3 cm) charged with either 100 µl of juice (grape or noni) or 100 µl of mashed-ripe noni fruit and fixed on a metal holder. The fly tube was placed within the centre of the airstream and 85 cm downwind of the odour source. Flies leaving the tube and arriving on the landing platform within the first 10 min after release were counted.

### Olfactory trap assay

The two-choice olfactory trap assay was performed essentially as described^15^. For each experiment, the two traps contained either 300 µl noni (Nu3 GmbH) or grape juice (Beutelsbacher Fruchtsaftkelterei). When using noni and grape fruits as stimuli, ripe fruits were homogenized with pestles and each trap filled with a spatula of the mix (to a volume equivalent to ∼300 µl juice). 25 fed, mated, ice-anesthetised female flies (3-5 days post-eclosion) were used for each experiment. The distribution of flies was scored after 24 h at 25°C under red light at 60% relative humidity; experiments with >25% dead flies in the arena after 24 h were discarded. The attraction index was calculated as follows: (number of flies in noni juice trap - number of flies in grape juice trap)/number of flies alive.

### Two-photon calcium imaging

Flies were mounted and dissected as previously described^71^, and images were acquired using a commercial upright 2-photon microscope (Zeiss LSM 710 NLO). In detail, an upright Zeiss AxioExaminer Z1 was fitted with a Ti:Sapphire Chameleon Ultra II infrared laser (Coherent) as excitation source. Images were acquired with a 20x water dipping objective (Plan-Apochromat 20x W; NA 1.0), with a resolution of 128×128 pixels (1.1902 pixels µm^−1^) and a scan speed of 12.6 µs pixel^−1^. Excitation wavelength was set to 920 nm at a laser output of 64.1-70.2 mW measured at the exit of the objective. Emitted light was filtered with a 500-550 nm band-pass filter, and photons were collected by an external non-descanned detector. Each measurement consisted of 50 images acquired at 4.13 Hz, with stimulation starting ∼5 s after the beginning of the acquisition and lasting for 1 s. Fly antennae were stimulated using a custom-made olfactometer as previously described^72^ with minor modifications. In brief, the fly antenna was permanently exposed to air flowing at a rate of 1.5 l min^−1^ and with 55% relative humidity obtained by combining a main stream of humidified room air (0.5 l min^−1^) and a secondary stream (1 l min^−1^) of normal room air. Both air streams were generated by vacuum pumps (KNF Neuberger AG) and the flow rate was controlled by two independent rotameters (Analyt). The secondary air stream was guided either through an empty 2 ml syringe or through a 2 ml syringe containing 20 µl of odour or solvent on a small cellulose pad (Kettenbach GmbH) to generate 1 s odour pulses. To switch between control air and odour stimulus application, a three-way magnetic valve (The Lee Company, Westbrook, CT) was controlled using Matlab via a VC6 valve controller unit (Harvard Apparatus). Data were processed using Fiji^73^ and custom written programs in Matlab and R as previously described^72^. Since bleaching was very strong at the beginning of each acquisition, the first 1.5 s were not considered for the analysis, therefore bleach correction was not required. Colour-coded images and boxplots show the peak response calculated as the mean relative change in fluorescence (% ΔF/F) of three frames around the maximum between frames 19 and 30.

### Wide-field calcium imaging

Flies were mounted and dissected as previously described^71^. Images were acquired with a CCD camera (CoolSNAP-HQ2 Digital Camera System) mounted on a fluorescence microscope (upright fixed stage Carl Zeiss Axio Examiner D1) equipped with a 40x water-immersion objective (W “Plan-Apochromat” 40x/1,0 VIS-IR DIC). Excitation light of 470 nm was produced with an LED light (Cool LED pE-100, VisiChrome). Light was guided through a filter block consisting of a 450– 490nm excitation filter, a dichroic mirror (T495LP), and a 500–550 nm emission filter (Chroma ET). Binned image size was 266×200 pixels on the chip, corresponding to 149×112 μm in the preparation. Exposure time varied between 80 and 100 ms to adjust for different basal fluorescence values across preparations. Films (12.5 s duration) were recorded with an acquisition rate of 4 Hz. Metafluor software (Visitron) was used to control the camera, light, data acquisition and onset of odour stimulation. Odour stimulation and data analysis were otherwise performed as described for two-photon calcium imaging.

### Electrophysiology

Single sensillum electrophysiological recordings were performed as described previously^74^. Noni and grape juice were purchased from Nu3 (nu3 GmbH) and Beutelsbacher (Beutelsbacher Fruchtsaftkelterei GmbH) and chemicals of the highest purity available from Sigma-Aldrich. Odorants were used at 1% (v/v) in all experiments unless noted otherwise in the figures or figure legends. Solvents were either double-distilled water (for noni juice, butyric acid (CAS 107-92-6), hexanoic acid (CAS 1821-02-9)) or paraffin oil (for octanoic acid (CAS 124-07-2), methyl butanoate (CAS 623-42-7), methyl hexanoate (CAS 106-70-7), methyl octanoate (CAS 111-11-5), ethyl butanoate (CAS 105-54-4), ethyl hexanoate (CAS 123-66-0), ethyl octanoate (CAS 106-32-1), *2*-heptanone (CAS 110-43-0), *1*-hexanol (CAS 111-27-3), and *3*-buten-*1*-ol (CAS 627-27-0)). Corrected responses were calculated as the number of spikes in a 500 ms window at stimulus onset (200 ms after stimulus delivery due to a delay by the air path) subtracting the number of spontaneous spikes in a 500 ms window 2 s before stimulation, multiplied by two to obtain spikes s^−1^. The solvent-corrected responses shown in the figures were calculated by subtracting from the response to each diluted odour, the response obtained when stimulating with the corresponding solvent. Spike counts for all experiments are provided in Supplementary Table 7.

### Immunohistochemistry

Fluorescent RNA *in situ* hybridisation using digoxigenin- or fluorescein-labelled RNA probes and immunofluorescence on whole-mount antennae were performed essentially as described^72,75^. *D. sechellia* probe templates were generated by amplification of regions of genomic DNA (DSSC 14021-0248.07) using primer pairs listed in Supplementary Table 6; these were cloned into *pCR-Blunt II-TOPO* and sequenced. *D. sechellia* OR22a antibodies were raised in rabbits against the peptide epitope PHISKKPLSERVKSRD (amino acids 7-22), affinity-purified (Proteintech Groups, Inc) and diluted 1:250. Other antibodies used were: rabbit α-IR75a (RRID: AB_2631091) 1:100^76^, guinea pig α-IR75b (RRID: AB_2631093) 1:200^76^, rabbit α-IR64a 1:100^41^, rabbit α-ORCO 1:200^38^, guinea pig α-IR8a 1:500^37^, rabbit α-IR25a^77^, rabbit α-GFP 1:500 (Invitrogen). Alexa488- and Cy5-conjugated goat α-guinea pig IgG and goat α-rabbit IgG secondary antibodies (Molecular Probes; Jackson Immunoresearch) were used at 1:500. Immunofluorescence on adult brains was performed as described^78^ using mouse monoclonal antibody nc82 1:10 (Developmental Studies Hybridoma Bank), rat monoclonal α-Elav 1:10 (Developmental Studies Hybridoma Bank) and rabbit α-GFP 1:500 (Invitrogen).

### *D. sechellia* reference brain

*D. sechellia* (DSSC 14021-0248.07) brains (2-7 day old animals) were stained with nc82 and imaged as described^79^. From 88 central female brains imaged, 26 high-quality confocal stacks were selected for averaging on a selected “seed” brain, essentially as described^80,81^. Similarly, a male reference brain (not shown here) was constructed, using 20 high-quality confocal stacks (from 87 initially imaged). Reciprocal bridging registrations between *D. melanogaster* and *D. sechellia* references brains were also generated to permit comparison of homologous neurons within a common template, essentially as described^81^. The reference brains (*DsecF* and *DsecM*), bridging registrations and associated code are available for download via http://jefferislab.org/si/auer2019.

### Image acquisition and processing

Confocal images of antennae and brains were acquired on an inverted confocal microscope (Zeiss LSM 710) equipped with an oil immersion 40x objective (Plan Neofluar 40x Oil immersion DIC objective; 1.3 NA) unless stated otherwise. Images were processed in Fiji^73^. *D. sechellia* brains were imaged and registered to a *D. sechellia* reference brain using the Fiji CMTK plugin (https://github.com/jefferis/fiji-cmtk-gui) as described^82^. For segmentation of individual glomeruli of the antennal lobe, glomerular identity was confirmed by location and labelling with *Gal4* reporters (Supplementary Fig. 2) and segmentation performed using Amira 6.5 (ThermoScientific). Glomerular volumes were calculated following segmentation with the Segmentation Editor plugin of Fiji using the 3D Manager Plugin. Olfactory sensory neuron numbers were counted using the Cell Counter Plugin in Fiji or Imaris (Bitplane). Projection neuron morphologies were reconstructed and measured in neuTube 1.0z^83^.

### Labelling of projection neurons

For photoactivation experiments we generated transgenic *D. sechellia* flies bearing one copy each of *nSyb-Gal4* and *UAS-C3PAGFP*^48^; *D. melanogaster* flies carried two copies of *UAS-C3PAGFP*, one copy of *UAS-SPAGFP*^48^ and one copy of *nSyb-Gal4*. Photoactivation was performed as described^48^ on adult female flies aged 3-5 days after eclosion. Brains were dissected in saline^84^ low carbonate (2 mm Mg^2+^ pH 7.2) and treated with collagenase (2 mg/ml, 45 s). We initially imaged dissected brains at 925 nm to identify the DM2 glomerulus based on anatomical position. Photoactivation was achieved through multiple cycles of exposure to 710-nm laser light with 15 min rest period between each photoactivation cycle to allow diffusion of the photoactivated fluorophore. Photoactivation and imaging was performed on an Ultima two-photon laser scanning microscope (Bruker) equipped with galvanometers driving a Chameleon XR laser (Coherent). Emitted photons were collected with a GaAsP photodiode detector (Bruker) or a PMT detector through a 60X objective (Olympus 60X water immersion; 0.9 NA).

PN dye-fillings were performed as described^48^ with some modifications. In brief, brains were dissected in saline, briefly treated with collagenase (2 mg ml^−1^, 45 s), washed and pinned with fine tungsten wires to a Sylgard sheet (World Precision Instruments) in a 35 mm Petri dish (Falcon) filled with saline. Pulled glass electrodes were backfilled with Texas Red Dextran (3000, lysine fixable, Thermo Scientific). The electrode was targeted to the DM2 glomerulus and the dye electroporated by applying voltage pulses (30 V) until it became visible in distal neural processes of the PN. The dye was left to diffuse for 60 min and brains subsequently imaged by two-photon microscopy as described above.

To label single PNs, the DM2 glomerulus was first subjected to one cycle of exposure to 710-nm laser light to identify the cell bodies of DM2 PNs. Subsequently, the filled glass electrode was placed in the centre of the soma of one DM2 PN and the dye was electroporated by applying voltage pulses (30 V) until it became visible in distal neural processes of the PN. The dye was left to diffuse for 60 min, the brains were fixed with 2% paraformaldehyde for 45 minutes and subjected to an antibody staining using nc82 (1:20) and anti-mouse Alexa Fluor 488 (1:500) as described above. Images were acquired on a Zeiss 880 Airy scan confocal microscope using a 40X objective (Plan Neofluar 40X oil immersion DIC objective; 1.3 NA).

### Statistics and reproducibility

Data were analysed and plotted using Excel and R (v3.2.3; R Foundation for Statistical Computing, Vienna, Austria, 2005; R-project-org) (code available upon request).

### Data availability

All relevant data supporting the findings of this study are available from the corresponding authors on request.

## Supplementary Figures and Table Legends

**Supplementary Figure 1.**
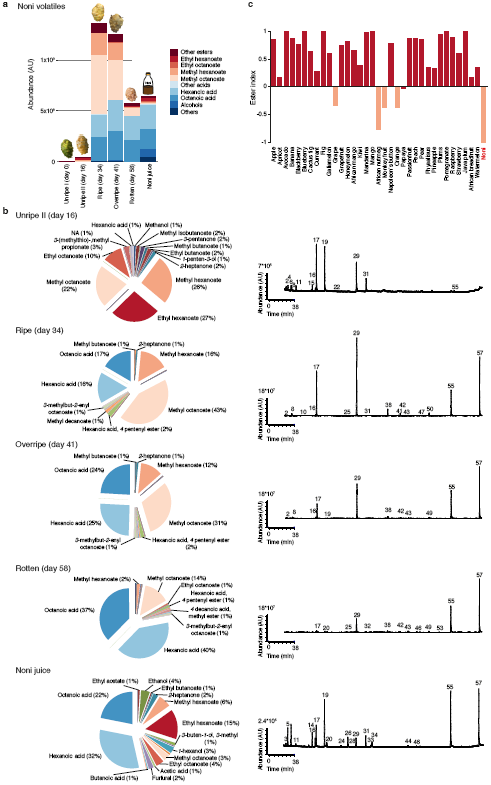
Volatile chemicals emitted by noni fruit and juice. **a**, Principal constituents of the odour bouquet of noni fruit at different ripening stages and commercial noni juice as determined by gas chromatography/mass spectrometry. AU = arbitrary units. **b**, Chemical composition of the odour bouquet of noni fruit at different stages of ripening and noni juice. Representative gas chromatograms are shown on the right. Numbers correspond to compounds as listed in Supplementary Table 1 (not all identified peaks are shown). NA = no specific compound could be assigned. **c**, Relative amount of ethyl- and methyl esters in 35 different fruits from data reported in^85^ and this study. The ester index is calculated as follows: (amount of ethyl esters - amount of methyl esters)/amount of both.

**Supplementary Figure 2.**
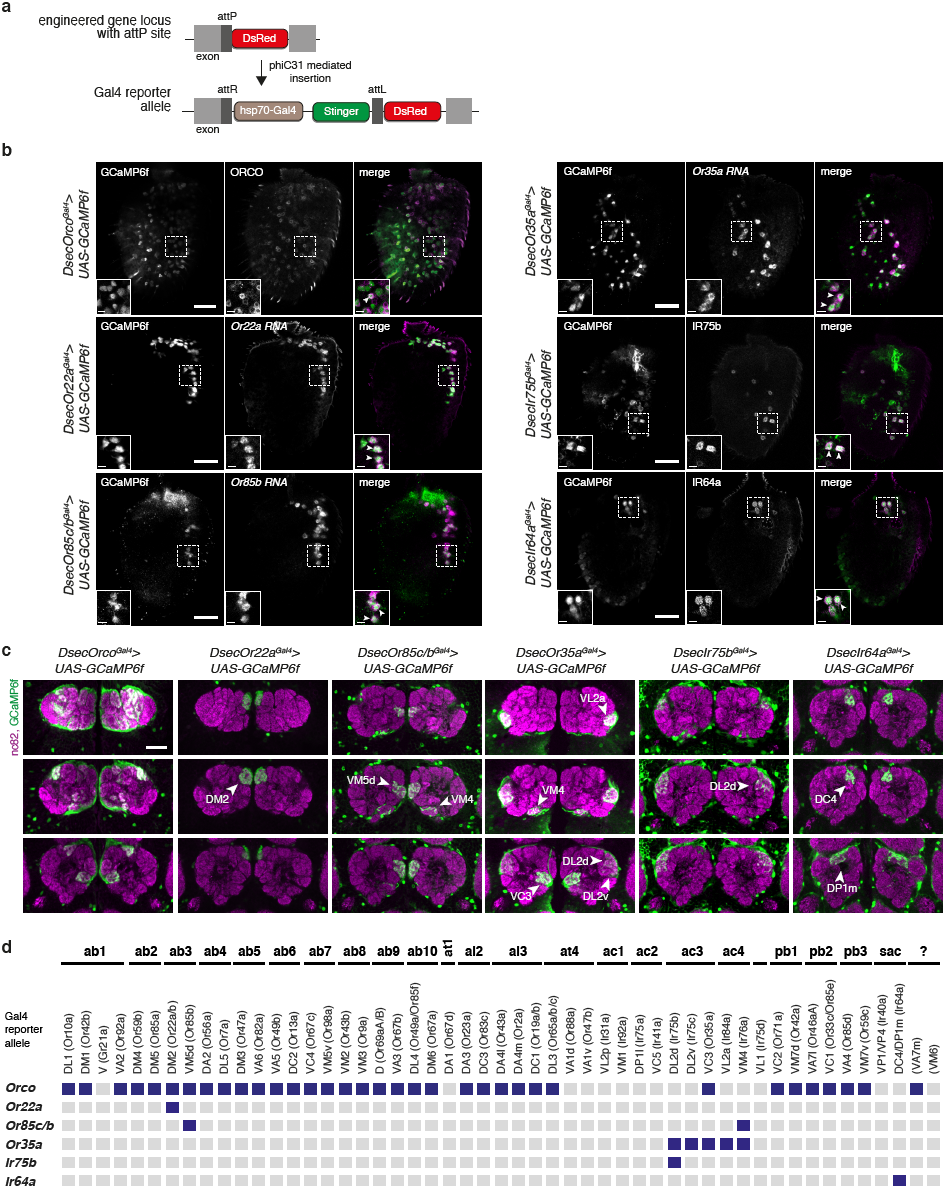
Olfactory sensory neuron Gal4 driver lines in *D. sechellia*. **a**, Schematic of the *Gal4* reporter allele generation strategy, through CRISPR/Cas9-mediated integration of an *attP* site (marked by *3xP3:DsRed*) into the desired *Or* or *Ir* locus (see Supplementary Fig. 4-5 for details), followed by introduction of a *Gal4* open reading frame via phiC31-mediated transgenesis. **b**, Co-expression of the indicated *Or*^*Gal4*^- or *Ir*^*Gal4*^-driven GCaMP6f signal (detected by α-GFP) with the corresponding receptor protein or transcript in whole-mount antennae. Arrowheads point towards examples of co-labelled cells. Scale bar = 25 µm; inset scale bar = 5 µm. **c**, Expression of the indicated *Or*^*Gal4*^- or *Ir*^*Gal4*^-driven GCaMP6f signal (detected by α-GFP) in glomeruli of the antennal lobe (the neuropil is visualised with nc82 (magenta); three focal planes are shown). Images were registered to a *D. sechellia* reference brain (see Methods) for better comparison of antennal lobe structure. Scale bar = 25 µm. **d**, Summary of the glomerular labelling by *Or*^*Gal4*^ or *Ir*^*Gal4*^ drivers as characterised in **c** (dark blue indicates GCaMP6f signal was detected). Glomeruli are organised by the compartmentalisation of the corresponding OSN populations in different sensilla classes (based on data in *D. melanogaster*^86^, ab: antennal basiconic; at: antennal trichoid; ai; antennal intermediate; ac: antennal coeloconic, pb = palp basiconic, sac = sacculus, ? = OSN population unknown). *Orco*^*Gal4*^ is expressed in most but not all (e.g., Or67d/DA1) expected OSN populations; *Or35a*^*Gal4*^ and *Or85c/b*^*Gal4*^ display some ectopic expression, reflecting in their labelling of >1 glomerulus.

**Supplementary Figure 3.**
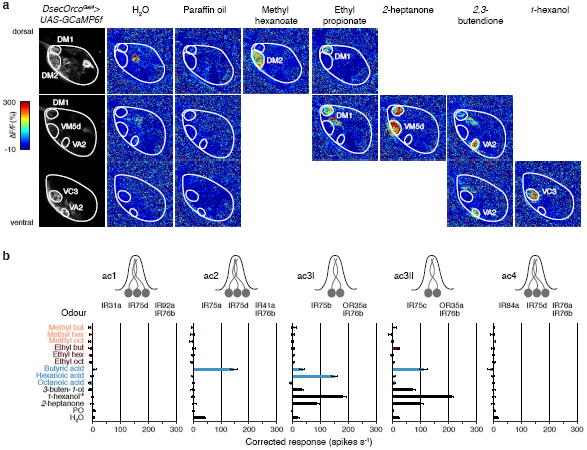
Physiological characterisation of noni-responsive olfactory pathways. **a**, Representative odour-evoked calcium responses in the axon termini of Orco OSNs in the *D. sechellia* antennal lobe (genotype: *UAS-GCaMP6f/UAS-GCaMP6f;;DsecOrco*^*Gal*^*4*^^*/+*) acquired by two-photon imaging. Three focal planes are shown, revealing different glomeruli along the dorsoventral axis. Left column: raw fluorescence images. Other columns: relative increase in GCaMP6f fluorescence (ΔF/F%) after stimulation with diagnostic odours. Methyl hexanoate (10^−6^) was used as diagnostic odour for Or22a/DM2, ethyl propionate (10^−2^) for Or42b/DM1, *2*-heptanone (10^−5^) for Or85b/VM5d, *2,3*-butendione (10^−2^) for Or92a/VA2 and *1*-hexanol (10^−4^) for Or35a/VC3. Glomerular boundaries are outlined. **b**, Electrophysiological responses in the antennal coeloconic (ac) sensilla classes to the indicated stimuli (mean ± SEM; n = 6-11, females) in *D. sechellia* (DSSC 14021-0248.07) representing the summed, solvent-corrected activities of the neurons they house, indicated in the cartoons on top.

**Supplementary Figure 4.**
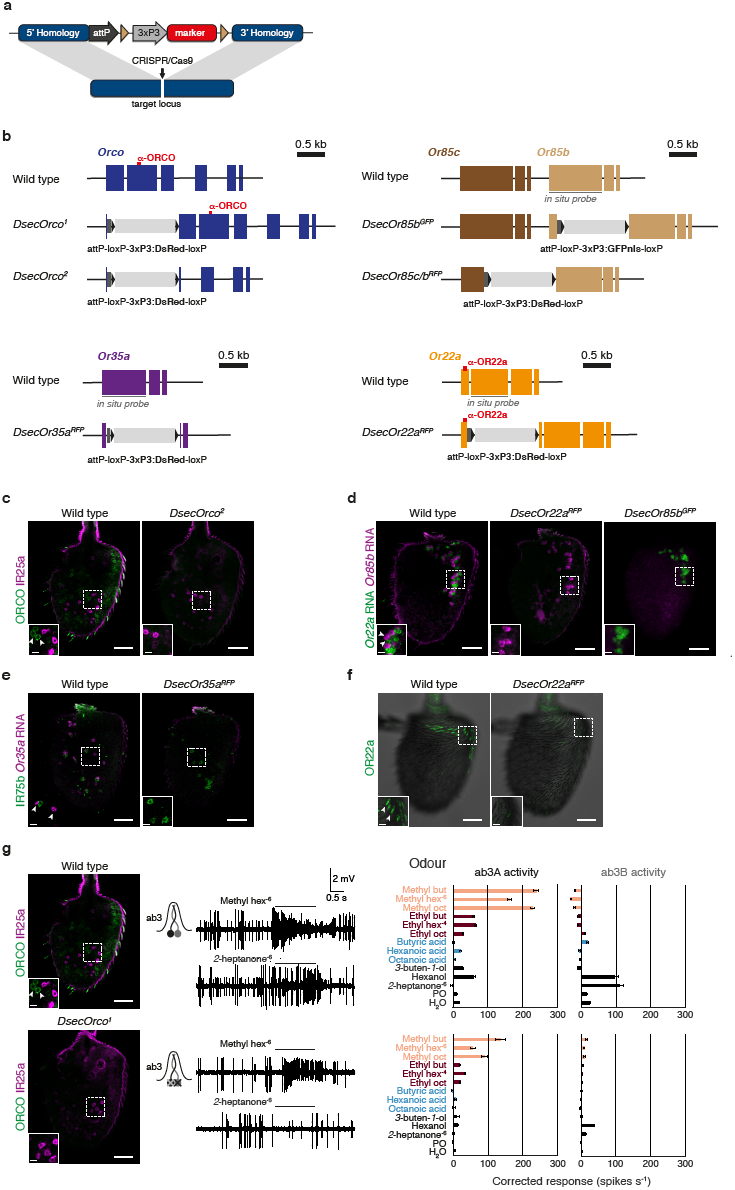
Generation and validation of loss-of-function alleles of *D. sechellia Or* genes. **a**, Schematic of the strategy for generating olfactory receptor mutant alleles, through integration of an eye-expressed fluorescent marker (*3xP3:DsRed* or *3xP3:GFPnls*) into the desired locus via CRISPR/Cas9-cleavage induced homologous recombination; integration was usually accompanied by deletion of one or more coding exons. Brown triangles: *loxP* sites for removal of the fluorescent marker via Cre recombination. **b**, Schematics depicting *Or* gene organisation and the structure of the mutant alleles. The location of the sequence encoding epitopes recognised by antibodies and regions corresponding to RNA probe templates are indicated. **c**, Immunostaining for ORCO and (as internal staining control) IR25a on whole-mount antennae from wild-type and *DsecOrco*^*^2^*^ animals. Arrowheads depict ORCO expressing cells. **c**-**g**,: Scale bar = 25 µm, inset scale bar = 5 µm. **d**, RNA FISH for *Or22a* and *Or85b* on whole-mount antennae from wild-type, *DsecOr22a*^*RFP*^ and *DsecOr85b*^*GFP*^ mutant animals. Arrowheads depict *Or22a* and *Or85b* expressing cells. **e**, Immunostaining for IR75b and RNA FISH for *Or35a* on whole-mount antennae from wild-type and *DsecOr35a*^*RFP*^ mutant animals. Arrowheads depict *Or35a* expressing cells. Note that *Or35a* neurons also pair with *Ir75c* neurons in ac3II sensilla^15^ leading to *Or35a* positive cells non-paired with IR75b in wild-type antenna. **f**, Immunostaining for OR22a on whole-mount antennae from wild-type and *DsecOr22a*^*RFP*^ mutant animals. Arrowheads depict OR22a housing sensilla. **g**, Left panel: Immunostaining for ORCO and (as internal staining control) IR25a on whole-mount antennae from wild-type (same picture as shown in **b**) and *DsecOrco*^*^1^*^ animals. Arrowheads depict ORCO expressing cells. Central panel: electrophysiological responses in the two neurons of the ab3 sensillum (compare Fig. 2) to odours present in noni (mean ± SEM; n = 5-11, females only) in *D. sechellia* (DSSC 14021-0248.07) and *DsecOrco*^*^1^*^ mutants. Representative response traces to methyl hexanoate (10^−6^) and *2*-heptanone (10^−6^) are shown to the left. Histograms represent the solvent-corrected activities per neuron. Surprisingly, even though ORCO expression is not detectable, weak electrophysiological responses in ab3 sensilla (and other ORCO-dependent sensilla (data not shown)) can be detected, suggesting trace levels of functional receptor protein are produced from this allele (which retains most coding exons). It is for this reason that we generated an independent, true null allele (*Orco*^*^2^*^) (Fig. 2a).

**Supplementary Figure 5.**
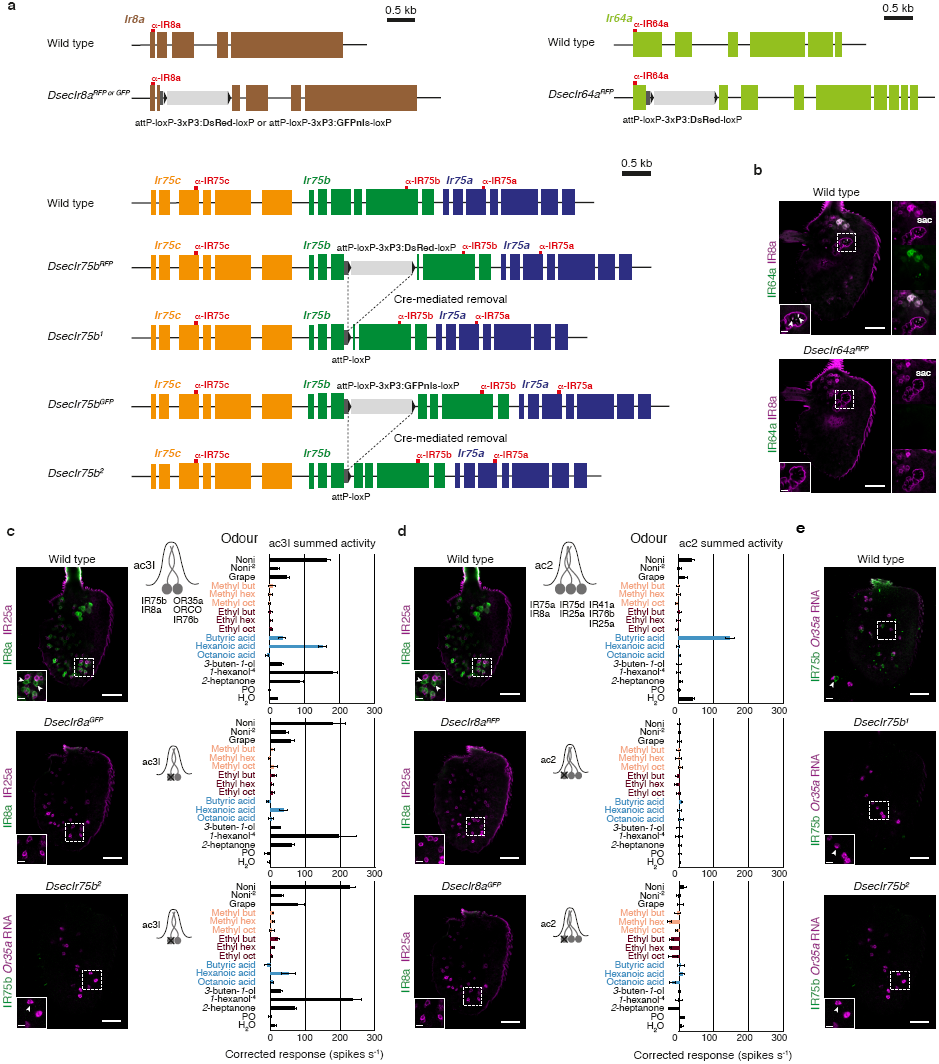
Generation and validation of loss-of-function alleles of *D. sechellia Ir* genes. **a**, Schematics depicting *Ir* gene organisation and the structure of the mutant alleles. For both alleles of *Ir75b*, the fluorophore was removed via Cre-mediated recombination. For all genes the sequence encoding epitopes used for immunofluorescence detection are indicated. **b**, Immunostaining for IR64a on whole-mount antennae from wild-type and *DsecIr64a*^*RFP*^ mutant animals. Arrowheads depict the sacculus innervating dendrites of Ir64a neurons. Scale bar = 25 µm, inset scale bar = 5 µm. **c**, Left panels: immunostaining for IR8a and (as internal staining control) IR25a on whole-mount antennae from wild-type and *DsecIr8a*^*GFP*^ animals (top, arrowheads depict IR8a expressing cells); immunostaining for IR75b and RNA FISH for *Or35a* on whole-mount antennae from *DsecIr75b*^*^2^*^ mutant animals (bottom, arrowhead depicts *Or35a* expressing cells). Scale bar = 25 µm, inset scale bar = 5 µm. Right panels: electrophysiological responses in the ac3I sensillum (neurons housed are indicated in the cartoon) to noni juice, grape juice and odours present in noni (mean ± SEM; n = 4-10, females) in *D. sechellia* (DSSC 14021-0248.07) and olfactory receptor mutants affecting the Ir75b neuron (*DsecIr8a*^*GFP*^, *DsecIr75b*^*^2^*^). Histograms represent the summed, solvent-corrected activities of the sensillum. **d**, Left panel: immunostaining for IR8a and (as internal staining control) IR25a on whole-mount antennae from wild-type (same picture as shown in **c**) and *DsecIr8a*^*RFP*^ and *DsecIr8a*^*GFP*^ animals. Arrowheads depict IR8a expressing cells. Scale bar = 25 µm, inset scale bar = 5 µm. Right panel: electrophysiological responses in the ac2 sensillum (neurons housed are indicated in the cartoon) to noni juice, grape juice and odours present in noni (mean ± SEM; n = 3-11, females only) in *D. sechellia* (DSSC 14021-0248.07) and olfactory receptor mutants affecting the Ir75a neuron (*DsecIr8a*^*RFP*^, *DsecIr8a*^*GFP*^). **e**, Immunostaining for IR75b and RNA FISH for *Or35a* on whole-mount antennae from wild-type and *DsecIr75b*^*^1^*^ and *DsecIr75b*^*^2^*^ mutant animals. Arrowheads depict *Or35a* expressing cells. Scale bar = 25 µm, inset scale bar = 5 µm.

**Supplementary Figure 6.**
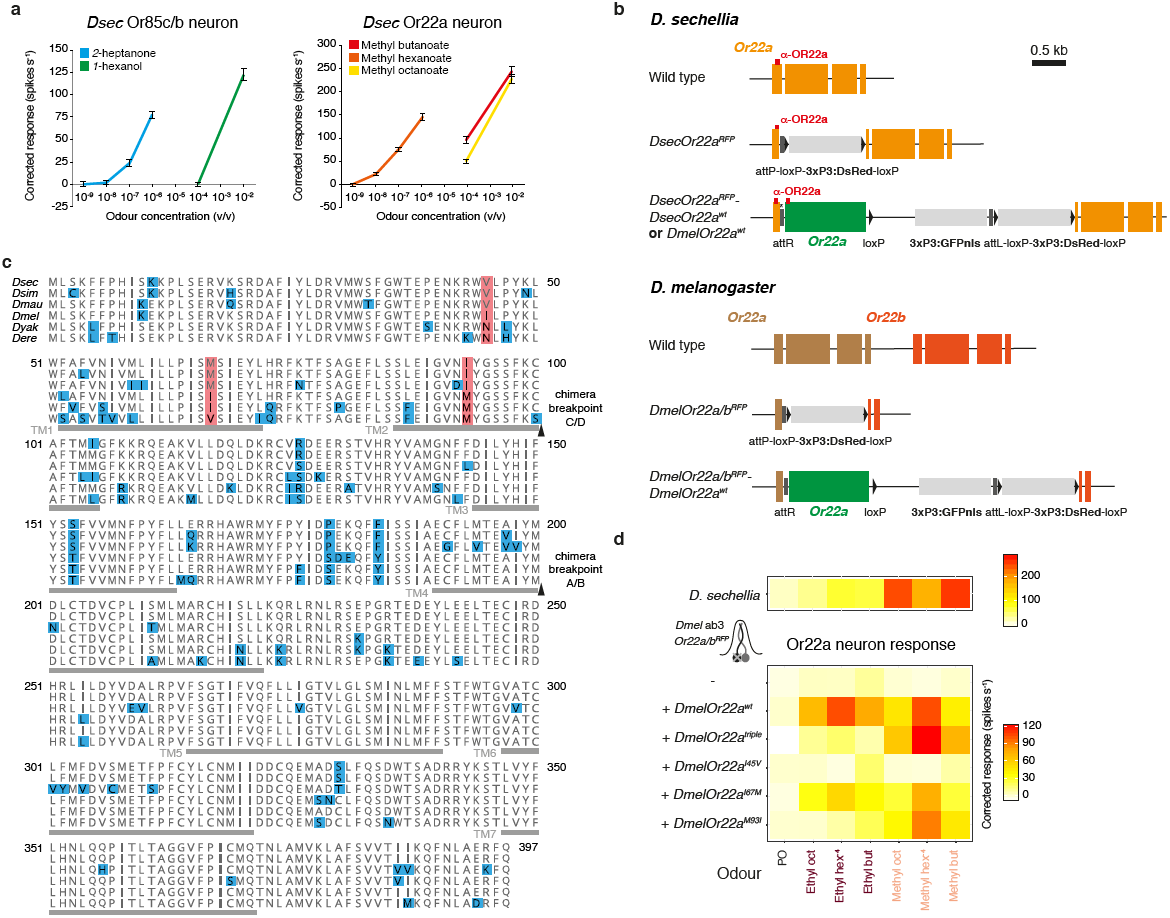
Ligand sensitivity of Or85c/b and Or22a neurons and genomic modifications at the *Or22a* locus. **a**, Left: electrophysiological responses of Or85c/b neurons (ab3B) upon stimulation with increasing concentrations of *2*-heptanone and *1*-hexanol in *D. sechellia* (mean ± SEM, n = 10-11, females). Right: electrophysiological responses of Or22a neurons (ab3A) upon stimulation with increasing concentrations of methyl butanoate, methyl hexanoate and methyl octanoate in *D. sechellia* (mean ± SEM, n = 10-11, females). **b**, Schematics depicting the arrangement of wild-type, mutant and rescue allele versions of the *DsecOr22a* gene (top) and *DmelOr22a/Or22b* (bottom). **c**, Protein alignment of OR22a orthologues of six species within the *D. melanogaster* species subgroup. Red shading = amino acid differences between *D. melanogaster* and *D. sechellia, D. simulans* and *D. mauritiana* (analysed by mutagenesis); blue shading = all other sequence differences. Arrowheads indicate chimera breakpoints (Fig. 4e). TM = transmembrane domain (location as in^13^). **d**, Physiological responses of OR22a variants expressed from the *Or22a/b* locus of *D. melanogaster* (n = 5-7, females). The location of each mutated residue is indicated in **c.** Upper row: *D. sechellia* wild-type response as shown in Fig. 4b.

**Supplementary Figure 7.**
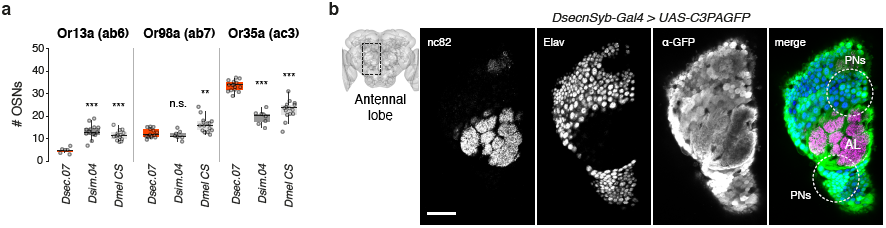
Peripheral and central olfactory circuit changes. **a**, Quantification of the number of OSNs expressing *Or13a* (ab6), *Or98a* (ab7) or *Or35a* (ac3I/II) in *D. sechellia* (DSSC 14021-0248.07), *D. simulans* (DSSC 14021-0251.004), and *D. melanogaster* (Canton-S) (n = 10-15, females). Comparisons to *Dsec.07* cell number counts are shown (pairwise Wilcoxon rank-sum test and *P* values adjusted for multiple comparisons using the Benjamini and Hochberg method): *** *P* < 0.001; ** *P* < 0.01; n.s. *P* > 0.05. **b**, Immunostaining with nc82 (neuropil), α-Elav (neurons) and α-GFP in a *DsecnSyb-Gal4/UAS-C3PAGFP* transgenic line, which expresses photoactivatable GFP pan-neuronally (genotype: *DsecnSypGal4/UAS-C3PAGFP;;)*. The schematic on the left shows the site of image acquisition in the brain. An anterior section through the antennal lobe (AL) is shown to reveal the position of the labelled projection neuron (PN) cell bodies (circled in the right panel). Scale bar = 25 µm.

**Supplementary Figure 8.**
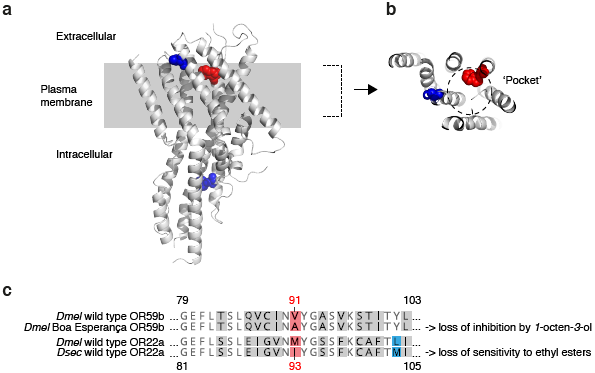
Location and conservation of ligand-binding determinants in *D. melanogaster* ORs. **a**, Side view of the ORCO monomer structure (determined by cryo-electronic microscopy^50^); the approximate location of the plasma membrane is indicated. The location of the corresponding residues of OR22a (based on alignments generated in^50^) are highlighted as spheres. Red = equivalent to position 93 in OR22a; blue = equivalent to position 45 and 67 in OR22a. **b**, A cross-section through the putative ligand-binding pocket of the ORCO structure shown in **a**. **c**, Partial protein sequence alignment of OR22a and OR59b. The equivalent residue to *D. melanogaster* OR22a M93 in OR59b (V91) exhibits intraspecific sequence variation, which influences odour sensitivity^51^.

**Supplementary Figure 9.**
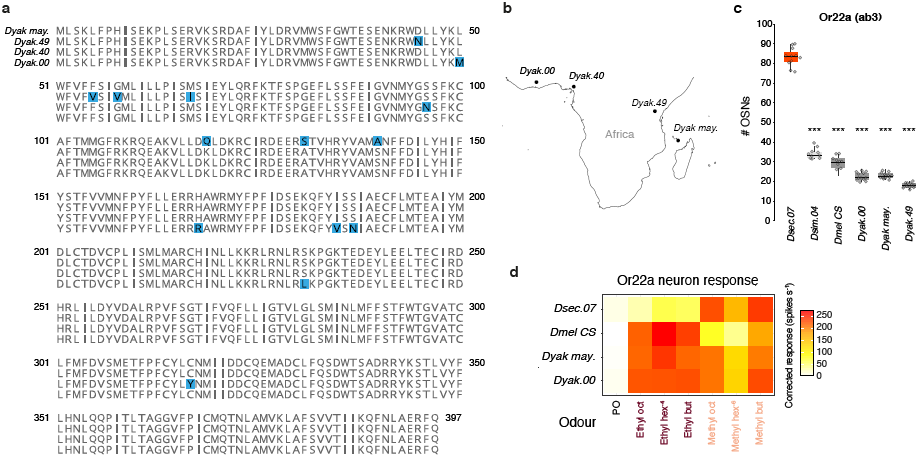
Analysis of Or22a neurons in *D. yakuba mayottensis*. **a**, Protein sequence alignment of OR22a sequences of four different *D. yakuba* strains: *D. yakuba* (DSSC 14021-0261.00, 14021-0261.40, 14021-0261.49) and *D. yakuba mayottensis*^87^ (*Dyak may.*). Blue shading highlights differences in these sequences. **b**, Collection sites of *D. yakuba* strains shown in **a**. DSSC 14021-0261.00: Ivory Coast; 14021-0261.40: Nguti, Cameroon; 14021-0261.49: Nairobi, Kenya; *D. yakuba mayottensis*: Mayotte. **c**, Quantification of the number of OSNs expressing *Or22a* RNA in *D. sechellia* (DSSC 14021-0248.07), *D. simulans* (DSSC 14021-0251.004), *D. melanogaster* (Canton-S) (data as shown in Fig. 5b), *D. yakuba* (DSSC 14021-0261.00, 14021-0261.49) and *D. yakuba mayottensis* (n = 10-12, females). Comparisons to *Dsec.07* cell number counts are shown (pairwise Wilcoxon rank-sum test and *P* values adjusted for multiple comparisons using the Benjamini and Hochberg method): *** *P* < 0.001. **d**, Physiological responses in the Or22a neurons of *D. sechellia* (DSSC 14021-0248.07), *D. melanogaster* (Canton-S) (data as shown in Fig. 4b), *D. yakuba* (DSSC 14021-0261.00) and *D. yakuba mayottensis* (n = 5-11, females).

**Supplementary Table 1. Gas Chromatography coupled Mass-Spectrometry analysis of noni fruit stages and fruit juices.**

Volatile compounds in five maturation stages of noni fruit and noni and grape juice (3 replicates/sample; values represent the mean (+/-standard deviation) of the peak area covered). In total, 57 compounds (arranged according to their retention times) have been identified with high probability based on matches to the NIST (National Institute of Standards and Technology) library. Compound names in blue: juice-specific compounds, in orange: fruit-specific compounds, in black: present in fruit and juice. Seventeen compounds marked with asterisk were newly identified in our study compared to^88,89^.

*(provided as separate Excel file)*

**Supplementary Table 2.**
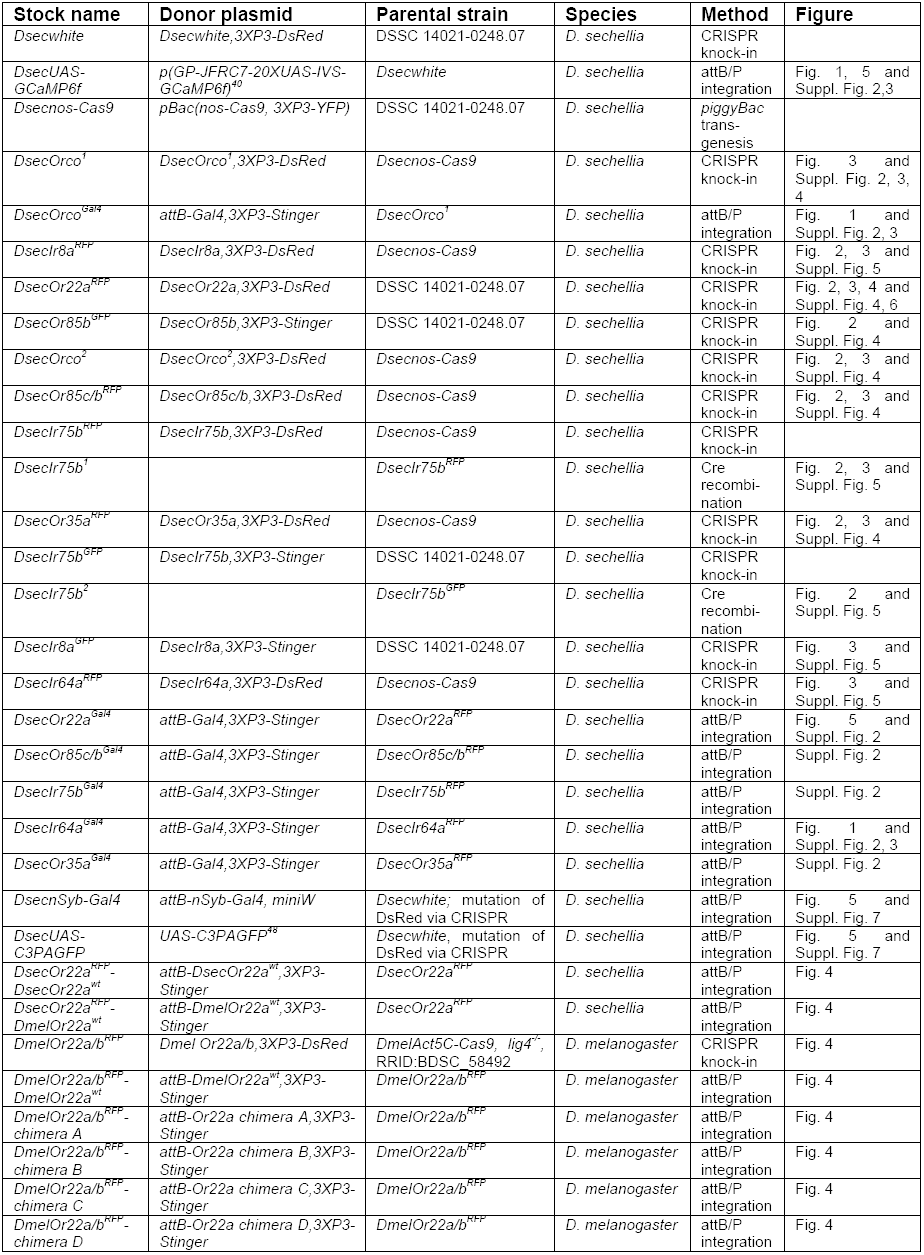
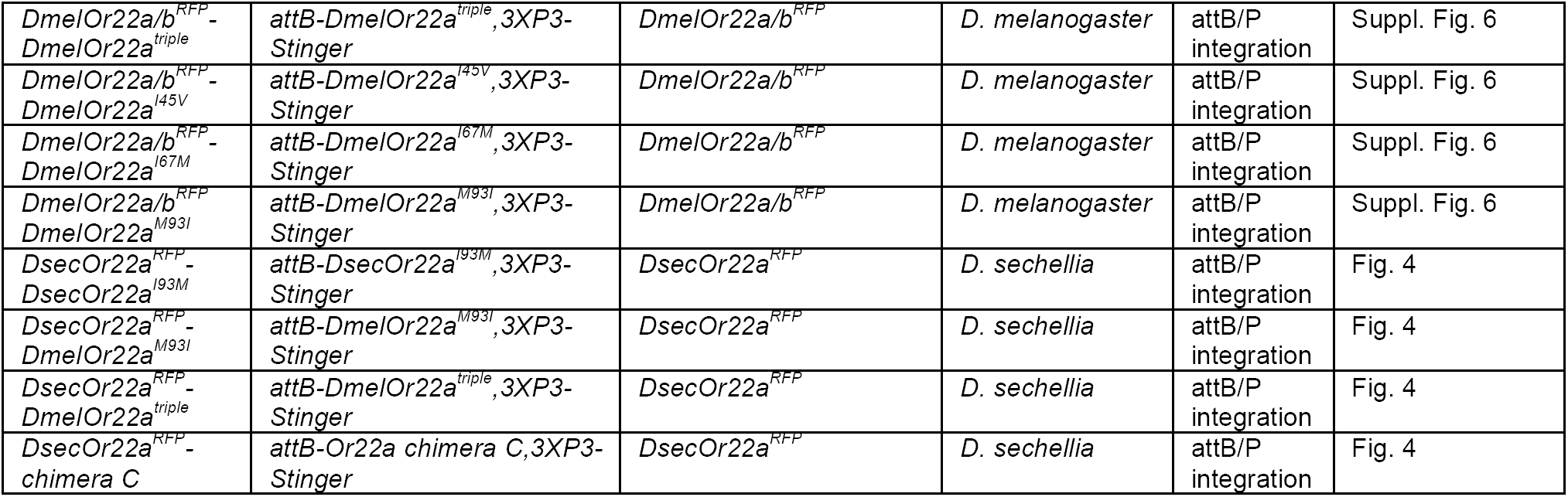
Details of the transgenic lines generated in this study.

**Supplementary Table 3.**
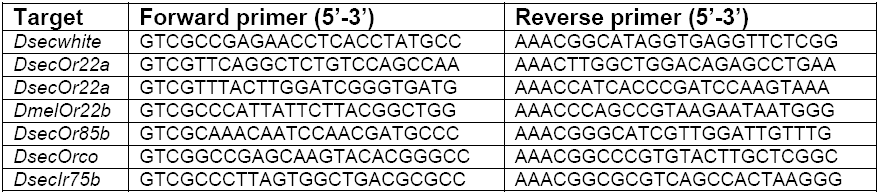
Oligonucleotides used to generate single sgRNA expression vectors.

**Supplementary Table 4. Oligonucleotides used to generate multi-sgRNA expression vectors.**

*(provided as a separate Excel file)*

**Supplementary Table 5.**
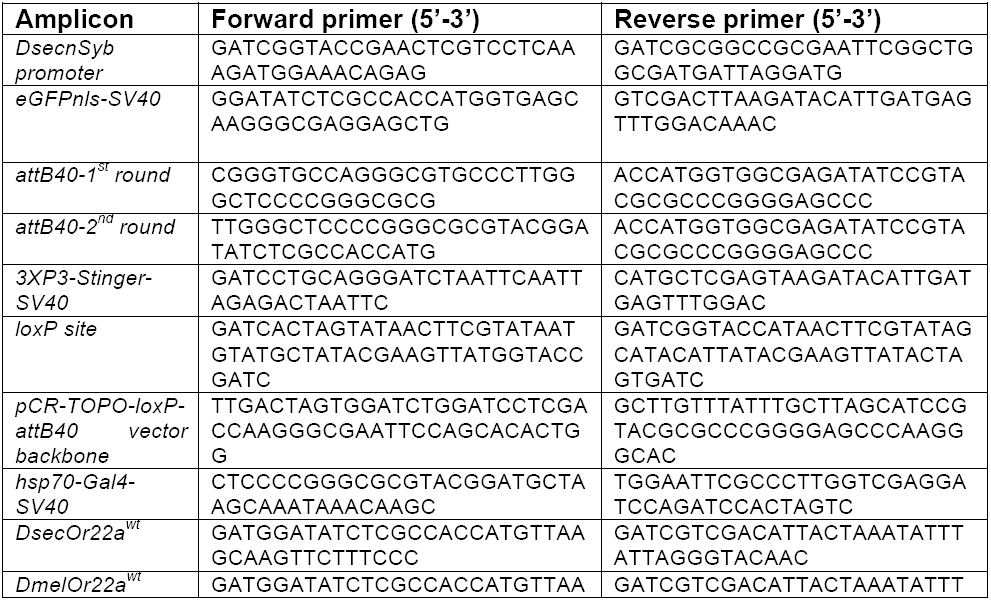
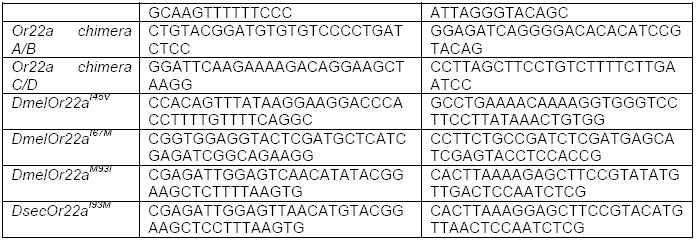
Oligonucleotides used for plasmid cloning and sitedirected mutagenesis.

**Supplementary Table 6.**
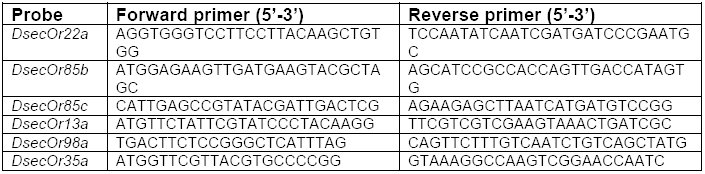
Oligonucleotides used to amplify RNA FISH probe templates.

**Supplementary Table 7. Spike counts for electrophysiological experiments.**

*(provided as a separate Excel file)*

## References

1 Arguello, J. R. & Benton, R. Open questions: Tackling Darwin’s “instincts”: the genetic basis of behavioral evolution. BMC Biol 15, 26, (2017).

2 Bendesky, A. & Bargmann, C. I. Genetic contributions to behavioural diversity at the gene-environment interface. Nat Rev Genet 12, 809–820, (2011).

3 Yalcin, B. et al. Genetic dissection of a behavioral quantitative trait locus shows that Rgs2 modulates anxiety in mice. Nat Genet 36, 1197–1202, (2004).

4 Osborne, K. A. et al. Natural behavior polymorphism due to a cGMP-dependent protein kinase of *Drosophila*. Science 277, 834–836, (1997).

5 Bendesky, A., Tsunozaki, M., Rockman, M. V., Kruglyak, L. & Bargmann, C. I. Catecholamine receptor polymorphisms affect decision-making in *C. elegans*. Nature 472, 313–318, (2011).

6 Bendesky, A. et al. The genetic basis of parental care evolution in monogamous mice. Nature 544, 434–439, (2017).

7 Weber, J. N., Peterson, B. K. & Hoekstra, H. E. Discrete genetic modules are responsible for complex burrow evolution in *Peromyscus* mice. Nature 493, 402–405, (2013).

8 Nozawa, M. & Nei, M. Evolutionary dynamics of olfactory receptor genes in *Drosophila* species. Proc Natl Acad Sci U S A 104, 7122–7127, (2007).

9 Markow, T. A. & O’Grady, P. Reproductive ecology of *Drosophila*. Functional Ecology 22, 747–759, (2008).

10 Crowley-Gall, A. et al. Population differences in olfaction accompany host shift in *Drosophila mojavensis*. Proc Biol Sci 283, (2016).

11 Linz, J. et al. Host plant-driven sensory specialization in *Drosophila erecta*. Proc Biol Sci 280, 20130626, (2013).

12 Keesey, I. W., Knaden, M. & Hansson, B. S. Olfactory specialization in *Drosophila suzukii* supports an ecological shift in host preference from rotten to fresh fruit. J Chem Ecol 41, 121–128, (2015).

13 Dekker, T., Ibba, I., Siju, K. P., Stensmyr, M. C. & Hansson, B. S. Olfactory shifts parallel superspecialism for toxic fruit in *Drosophila melanogaster* sibling, *D. sechellia*. Curr Biol 16, 101–109, (2006).

14 Ibba, I., Angioy, A. M., Hansson, B. S. & Dekker, T. Macroglomeruli for fruit odors change blend preference in *Drosophila*. Naturwissenschaften 97, 1059–1066, (2010).

15 Prieto-Godino, L. L. et al. Evolution of acid-sensing olfactory circuits in *Drosophilids*. Neuron 93, 661–676 e666, (2017).

16 Ding, Y., Berrocal, A., Morita, T., Longden, K. D. & Stern, D. L. Natural courtship song variation caused by an intronic retroelement in an ion channel gene. Nature 536, 329–332, (2016).

17 Seeholzer, L. F., Seppo, M., Stern, D. L. & Ruta, V. Evolution of a central neural circuit underlies *Drosophila* mate preferences. Nature 559, 564–569, (2018).

18 Lachaise, D. & Silvain, J. F. How two Afrotropical endemics made two cosmopolitan human commensals: the *Drosophila melanogaster* - *D.simulans* palaeogeographic riddle. Genetica 120, 17–39, (2004).

19 Garrigan, D. et al. Genome sequencing reveals complex speciation in the *Drosophila simulans* clade. Genome Res 22, 1499–1511, (2012).

20 Schrider, D. R., Ayroles, J., Matute, D. R. & Kern, A. D. Supervised machine learning reveals introgressed loci in the genomes of *Drosophila simulans* and *D. sechellia*. Plos Genetics 14, (2018).

21 Jones, C. D. The genetics of adaptation in *Drosophila sechellia*. Genetica 123, 137–145, (2005).

22 Stensmyr, M. C. *Drosophila sechellia* as a model in chemosensory neuroecology. Ann Ny Acad Sci 1170, 468–475, (2009).

23 R’Kha, S., Capy, P. & David, J. R. Host-plant specialization in the *Drosophila melanogaster* species complex: a physiological, behavioral, and genetical analysis. Proc Natl Acad Sci U S A 88, 1835–1839, (1991).

24 Lavista-Llanos, S. et al. Dopamine drives *Drosophila sechellia* adaptation to its toxic host. Elife 3, (2014).

25 Higa, I. & Fuyama, Y. Genetics of Food Preference in *Drosophila sechellia*. Responses to Food Attractants. Genetica 88, 129–136, (1993).

26 Matsuo, T., Sugaya, S., Yasukawa, J., Aigaki, T. & Fuyama, Y. Odorant-binding proteins OBP57d and OBP57e affect taste perception and host-plant preference in *Drosophila sechellia*. PLoS Biol 5, e118, (2007).

27 Cobb, M., Burnet, B., Blizard, R. & Jallon, J. M. Courtship in *Drosophila sechellia* - its structure, functional aspects, and relationship to those of other members of the *Drosophila melanogaster* species subgroup. J Insect Behav 2, 63–89, (1989).

28 Coyne, J. A. Genetics of sexual isolation in females of the *Drosophila simulans* species complex. Genet Res 60, 25–31, (1992).

29 Amlou, M., Moreteau, B. & David, J. R. Genetic analysis of *Drosophila sechellia* specialization: Oviposition behavior toward the major aliphatic acids of its host plant. Behav Genet 28, 455–464, (1998).

30 Moreteau, B., Rkha, S. & David, J. R. Genetics of a nonoptimal behavior - oviposition preference of *Drosophila mauritiana* for a toxic resource. Behav Genet 24, 433–441, (1994).

31 Erezyilmaz, D. F. & Stern, D. L. Pupariation site preference within and between *Drosophila* sibling species. Evolution 67, 2714–2727, (2013).

32 Jones, C. D. The genetic basis of *Drosophila sechellia’s* resistance to a host plant toxin. Genetics 149, 1899–1908, (1998).

33 Gleason, J. M. & Ritchie, M. G. Do quantitative trait loci (QTL) for a courtship song difference between *Drosophila simulans* and *D. sechellia* coincide with candidate genes and intraspecific QTL? Genetics 166, 1303–1311, (2004).

34 Dweck, H. K. et al. Olfactory channels associated with the *Drosophila* maxillary palp mediate short- and long-range attraction. Elife 5, (2016).

35 Vosshall, L. B. & Stocker, R. F. Molecular architecture of smell and taste in *Drosophila*. Annu Rev Neurosci 30, 505–533, (2007).

36 Larsson, M. C. et al. Or83b encodes a broadly expressed odorant receptor essential for *Drosophila* olfaction. Neuron 43, 703–714, (2004).

37 Abuin, L. et al. Functional architecture of olfactory ionotropic glutamate receptors. Neuron 69, 44–60, (2011).

38 Benton, R., Sachse, S., Michnick, S. W. & Vosshall, L. B. Atypical membrane topology and heteromeric function of *Drosophila* odorant receptors *in vivo*. PLoS Biol 4, e20, (2006).

39 Stensmyr, M. C., Dekker, T. & Hansson, B. S. Evolution of the olfactory code in the *Drosophila melanogaster* subgroup. Proc Biol Sci 270, 2333–2340, (2003).

40 Chen, T. W. et al. Ultrasensitive fluorescent proteins for imaging neuronal activity. Nature 499, 295–300, (2013).

41 Ai, M. et al. Acid sensing by the *Drosophila* olfactory system. Nature 468, 691–695, (2010).

42 Dobritsa, A. A., van der Goes van Naters, W., Warr, C. G., Steinbrecht, R. A. & Carlson, J. R. Integrating the molecular and cellular basis of odor coding in the *Drosophila* antenna. Neuron 37, 827–841, (2003).

43 Couto, A., Alenius, M. & Dickson, B. J. Molecular, anatomical, and functional organization of the *Drosophila* olfactory system. Curr Biol 15, 1535–1547, (2005).

44 Kreher, S. A., Kwon, J. Y. & Carlson, J. R. The molecular basis of odor coding in the *Drosophila* larva. Neuron 46, 445–456, (2005).

45 Shiao, M. S. et al. Expression divergence of chemosensory genes between *Drosophila sechellia* and its sibling species and its implications for host shift. Genome Biol Evol 7, 2843–2858, (2015).

46 Yao, C. A., Ignell, R. & Carlson, J. R. Chemosensory coding by neurons in the coeloconic sensilla of the *Drosophila* antenna. J Neurosci 25, 8359–8367, (2005).

47 Ai, M. et al. Ionotropic glutamate receptors IR64a and IR8a form a functional odorant receptor complex *in vivo* in *Drosophila*. J Neurosci 33, 10741–10749, (2013).

48 Ruta, V. et al. A dimorphic pheromone circuit in *Drosophila* from sensory input to descending output. Nature 468, 686–690, (2010).

49 Grabe, V. & Sachse, S. Fundamental principles of the olfactory code. Bio Systems 164, 94–101, (2018).

50 Butterwick, J. A. et al. Cryo-EM structure of the insect olfactory receptor ORCO. Nature 560, 447–452, (2018).

51 Pellegrino, M., Steinbach, N., Stensmyr, M. C., Hansson, B. S. & Vosshall, L. B. A natural polymorphism alters odour and DEET sensitivity in an insect odorant receptor. Nature 478, 511–514, (2011).

52 Aguade, M. Nucleotide and copy-number polymorphism at the odorant receptor genes *Or22a* and *Or22b* in *Drosophila melanogaster*. Mol Biol Evol 26, 61–70, (2009).

53 Goldman-Huertas, B. et al. Evolution of herbivory in *Drosophilidae* linked to loss of behaviors, antennal responses, odorant receptors, and ancestral diet. Proc Natl Acad Sci U S A 112, 3026–3031, (2015).

54 Guo, S. & Kim, J. Molecular evolution of *Drosophila* odorant receptor genes. Mol Biol Evol 24, 1198–1207, (2007).

55 de Bruyne, M., Smart, R., Zammit, E. & Warr, C. G. Functional and molecular evolution of olfactory neurons and receptors for aliphatic esters across the *Drosophila* genus. J Comp Physiol A Neuroethol Sens Neural Behav Physiol 196, 97–109, (2010).

56 Yassin, A. et al. Recurrent specialization on a toxic fruit in an island *Drosophila* population. Proc Natl Acad Sci U S A 113, 4771–4776, (2016).

57 Ambrose, D., Ellender, J. H., Lee, E. B., Sprake, C. H. S. & Townsend, R. Thermodynamic properties of organic oxygen compounds XXXVIII. Vapour pressures of some aliphatic ketones. The Journal of Chemical Thermodynamics 7, 453–472, (1975).

58 Daubert, T. E. & Danner, R. P. Physical and thermodynamic properties of pure chemicals data compilation. Book, (1997).

59 Port, F., Chen, H. M., Lee, T. & Bullock, S. L. Optimized CRISPR/Cas tools for efficient germline and somatic genome engineering in *Drosophila*. Proc Natl Acad Sci U S A, (2014).

60 Port, F. & Bullock, S. L. Augmenting CRISPR applications in *Drosophila* with tRNA-flanked sgRNAs. Nat Methods 13, 852–854, (2016).

61 Barolo, S., Carver, L. A. & Posakony, J. W. GFP and beta-galactosidase transformation vectors for promoter/enhancer analysis in *Drosophila*. Biotechniques 29, 726, 728, 730, 732, (2000).

62 Gratz, S. J. et al. Highly specific and efficient CRISPR/Cas9-catalyzed homology-directed repair in *Drosophila*. Genetics 196, 961–971, (2014).

63 Stern, D. L. Tagmentation-Based Mapping (TagMap) of Mobile DNA Genomic Insertion Sites. bioRxiv, (2016).

64 Croset, V. et al. Ancient protostome origin of chemosensory ionotropic glutamate receptors and the evolution of insect taste and olfaction. PLoS Genet 6, e1001064, (2010).

65 Bateman, J. R. & Wu, C. T. A simple polymerase chain reaction-based method for the construction of recombinase-mediated cassette exchange donor vectors. Genetics 180, 1763–1766, (2008).

66 Groth, A. C., Olivares, E. C., Thyagarajan, B. & Calos, M. P. A phage integrase directs efficient site-specific integration in human cells. Proc Natl Acad Sci U S A 97, 5995–6000, (2000).

67 Arnoult, L. et al. Emergence and diversification of fly pigmentation through evolution of a gene regulatory module. Science 339, 1423–1426, (2013).

68 Gratz, S. J. et al. Genome engineering of *Drosophila* with the CRISPR RNA-guided Cas9 nuclease. Genetics 194, 1029–1035, (2013).

69 Zhang, X., Koolhaas, W. H. & Schnorrer, F. A versatile two-step CRISPR- and RMCE-based strategy for efficient genome engineering in *Drosophila*. G3 (Bethesda) 4, 2409–2418, (2014).

70 Gohl, D. M. et al. A versatile *in vivo* system for directed dissection of gene expression patterns. Nat Methods 8, 231–237, (2011).

71 Knecht, Z. A. et al. Distinct combinations of variant ionotropic glutamate receptors mediate thermosensation and hygrosensation in *Drosophila*. Elife 5, (2016).

72 Silbering, A. F. et al. Complementary function and integrated wiring of the evolutionarily distinct *Drosophila* olfactory subsystems. J Neurosci 31, 13357–13375, (2011).

73 Schindelin, J. et al. Fiji: an open-source platform for biological-image analysis. Nat Methods 9, 676–682, (2012).

74 Benton, R. & Dahanukar, A. Electrophysiological recording from *Drosophila* olfactory sensilla. Cold Spring Harb Protoc 2011, 824–838, (2011).

75 Saina, M. & Benton, R. Visualizing olfactory receptor expression and localization in *Drosophila*. Methods Mol Biol 1003, 211–228, (2013).

76 Prieto-Godino, L. L. et al. Olfactory receptor pseudo-pseudogenes. Nature 539, 93–97, (2016).

77 Benton, R., Vannice, K. S., Gomez-Diaz, C. & Vosshall, L. B. Variant ionotropic glutamate receptors as chemosensory receptors in *Drosophila*. Cell 136, 149–162, (2009).

78 Sanchez-Alcaniz, J. A., Zappia, G., Marion-Poll, F. & Benton, R. A mechanosensory receptor required for food texture detection in *Drosophila*. Nat Commun 8, 14192, (2017).

79 Ostrovsky, A., Cachero, S. & Jefferis, G. Clonal analysis of olfaction in *Drosophila*: immunochemistry and imaging of fly brains. Cold Spring Harb Protoc 2013, 342–346, (2013).

80 Jefferis, G. S. et al. Comprehensive maps of *Drosophila* higher olfactory centers: spatially segregated fruit and pheromone representation. Cell 128, 1187–1203, (2007).

81 Manton, J. D. et al. Combining genome-scale *Drosophila* 3D neuroanatomical data by bridging template brains. bioRxiv, (2014).

82 Cachero, S., Ostrovsky, A. D., Yu, J. Y., Dickson, B. J. & Jefferis, G. S. Sexual dimorphism in the fly brain. Curr Biol 20, 1589–1601, (2010).

83 Feng, L., Zhao, T. & Kim, J. neuTube 1.0: A New Design for Efficient Neuron Reconstruction Software Based on the SWC Format. eNeuro 2, (2015).

84 Caron, S. J., Ruta, V., Abbott, L. F. & Axel, R. Random convergence of olfactory inputs in the *Drosophila* mushroom body. Nature 497, 113–117, (2013).

85 Dweck, H. K. M. et al. The Olfactory Logic behind Fruit Odor Preferences in Larval and Adult *Drosophila*. Cell Rep 23, 2524–2531, (2018).

86 Grabe, V. et al. Elucidating the neuronal architecture of olfactory glomeruli in the *Drosophila* antennal lobe. Cell Rep 16, 3401–3413, (2016).

87 Yassin, A. *Drosophila yakuba mayottensis*, a new model for the study of incipient ecological speciation. Fly (Austin), 1–9, (2016).

88 Farine, J. P., Legal, L., Moreteau, B. & Le Quere, J. L. Volatile components of ripe fruits of *Morinda citrifolia* and their effects on *Drosophila*. Phytochemistry 41, 433–438, (1995).

89 Pino, J. A., Marquez, E., Quijano, C. E. & Castro, D. Volatile compounds in noni (*Morinda citrifolia L.*) at two ripening stages. Ciencia Tecnol Alime 30, 183–187, (2010).

